# Shifting Redox Reaction Equilibria on Demand Using an Orthogonal Redox Cofactor

**DOI:** 10.1101/2023.08.29.555398

**Authors:** Derek Aspacio, Yulai Zhang, Youtian Cui, Edward King, William B. Black, Sean Perea, Emma Luu, Justin B. Siegel, Han Li

**Author notes:** **Corresponding Authors Han Li** - Department of Chemical and Biomolecular Engineering, University of California, Irvine, Irvine, California 92697, United States; Department of Biomedical Engineering, University of California, Irvine, California 92697-3900, United States. Department of Biological Chemistry, University of California, Irvine, Irvine, California 92697-3900, United States. These authors contributed equally (Derek Aspacio, Yulai Zhang, Youtian Cui).

## Abstract

Natural metabolism relies on chemical compartmentalization of two redox cofactors, NAD^+^ and NADP^+^, to orchestrate life-essential redox reaction directions. However, in whole cells the reliance on these canonical cofactors limits flexible control of redox reaction direction as these reactions are permanently tied to catabolism or anabolism. In cell-free systems, NADP^+^ is too expensive in large scale. We have previously reported the use of nicotinamide mononucleotide, (NMN^+^) as a low-cost, noncanonical redox cofactor capable of specific electron delivery to diverse chemistries.

Here, we present Nox Ortho, an NMNH-specific water-forming oxidase, that completes the toolkit to modulate NMNH/NMN^+^ ratio. This work uncovers an enzyme design principle that succeeds in parallel engineering of six butanediol dehydrogenases as NMN(H)-orthogonal biocatalysts consistently with a 10^3^ – 10^6^ -fold cofactor specificity switch from NAD(P)^+^ to NMN^+^. We combine these to produce chiral-pure 2,3-butanediol (Bdo) isomers without interference from NAD(H) or NADP(H) in vitro and in *E. coli* cells. We establish that NMN(H) can be held at a distinct redox ratio on demand, decoupled from both NAD(H) and NADP(H) redox ratios in vitro and in vivo.

## Introduction

Biomanufacturing of renewable chemicals by engineered microbes^1,2^ or synthetic biochemistry^3^ has the potential to supplant many petroleum-derived chemical industries^4^. However, many engineered biosynthetic pathways fail to proceed past laboratory scale, because of insufficient titer, productivity, or yield^5^. One prevalent reason is because the chassis organisms that provide high energy cofactors^6^ also contain numerous competing pathways, which drain these critical resources. We and others have proposed that orthogonal metabolic systems could solve this limitation through specific delivery of resources^7–11^.

Nature uses two different redox cofactors, nicotinamide adenine dinucleotide (NAD^+^) and nicotinamide adenine dinucleotide phosphate (NADP^+^), to specifically allocate electrons to catabolism and anabolism, respectively. Furthermore, by maintaining a low NADH/NAD^+^ ratio and a high NADPH/NADP^+^ ratio^12,13^, cells use two redox cofactors to provide separate driving forces that ensure catabolism and anabolism proceed in opposite directions^14^. This principle has also been recapitulated in cell-free systems^12,15,16^.

However, in whole-cell biomanufacturing, dependence on NAD(P)/H permanently ties the reaction direction of a desired redox reaction to either catabolism or anabolism. This does not allow flexible control of reaction equilibrium. In cell-free biomanufacturing, NADP(H) is avoided due to its formidable cost and low stability^17,18^. This leaves NAD(H) as the sole electron carrier, which cannot support both oxidation and reduction simultaneously without forming a futile cycle^12^.

These limitations associated with relying on natural cofactors motivate the development of noncanonical cofactors^8,10,11,19–25^. Previously, we established a noncanonical redox cofactor, nicotinamide mononucleotide (NMN^+^), which operates in an orthogonal fashion to NAD(P)^+^ to precisely channel reducing power in *Escherichia coli* whole cells and crude lysates^8,11,26–28^. NMN^+^ is also a lower cost alternative to NAD(P)^+ 29,30^, and we have leveraged it to sustain industrially relevant total turnover numbers (TTNs) in cell-free reactions^8,27^.

Here, we develop an NMNH-specific oxidase (Nox Ortho) derived from *Lactobacillus lactis* water-forming NADH oxidase (*Ll* Nox) via a high-throughput, growth-based selection^26^. Together with the NMN^+^-specific glucose dehydrogenase (Gdh Ortho) we previously engineered^8^, this work completes the toolkit to modulate NMNH/NMN^+^ ratio. We establish that NMN(H) cofactor can be held at a distinct redox ratio on demand (ranging from 13 to 0.07 for NMNH/NMN^+^) that is decoupled from both NAD(H) and NADP(H) redox ratios in vitro and in vivo. By designing S-specific butanediol dehydrogenases (Bdhs) to specifically utilize NMN(H), we tap into this orthogonal driving force to produce chiral-pure 2,3-butanediol (Bdo) with high degree of completion, without interference by the NAD(P)/H reduction potentials in vitro and in vivo.

More importantly, this work also points out a path to readily broadening the applications of this unwavering redox driving force. The enzyme design principle discovered from directed evolution of Nox Ortho, namely, to restrict the cofactor binding pocket with hydrogen bonding, is consistently translated onto six Bdh enzymes to create NMN(H)-orthogonal catalysts. On different enzyme scaffolds (only 47.3% pairwise identity between the two Bdhs characterized below), this strategy consistently results in 1.0 × 10^3^ – 3.0 × 10^6^ -fold switch of cofactor specificity from NADP(H) or NAD(H) to NMN(H) relative to WT. Given the conserved positions of these mutations, we envision rapidly extending this design principle to other redox enzymes that catalyze key steps in biomanufacturing where precise equilibrium control is needed.

## Results and Discussion

### Design of Bdo Stereo-upgrading System as a Test Bed of Orthogonal Redox Driving Force

Bdo is an important bio-based chiral chemical with broad industrial applications such as synthetic rubbers^31,32^, fuels^31,32^, and pharmaceuticals^32^. Bdo exists as three stereoisomers (Fig. 1), (2S,3R)-meso-butanediol (m-Bdo), (2S,3S)-butanediol (SS-Bdo), and (2R,3R)-butanediol (RR-Bdo). The goal here is to convert m-Bdo to either SS-Bdo or RR-Bdo with high purity, which is a value-added process^33–35^.

**Figure 1.**
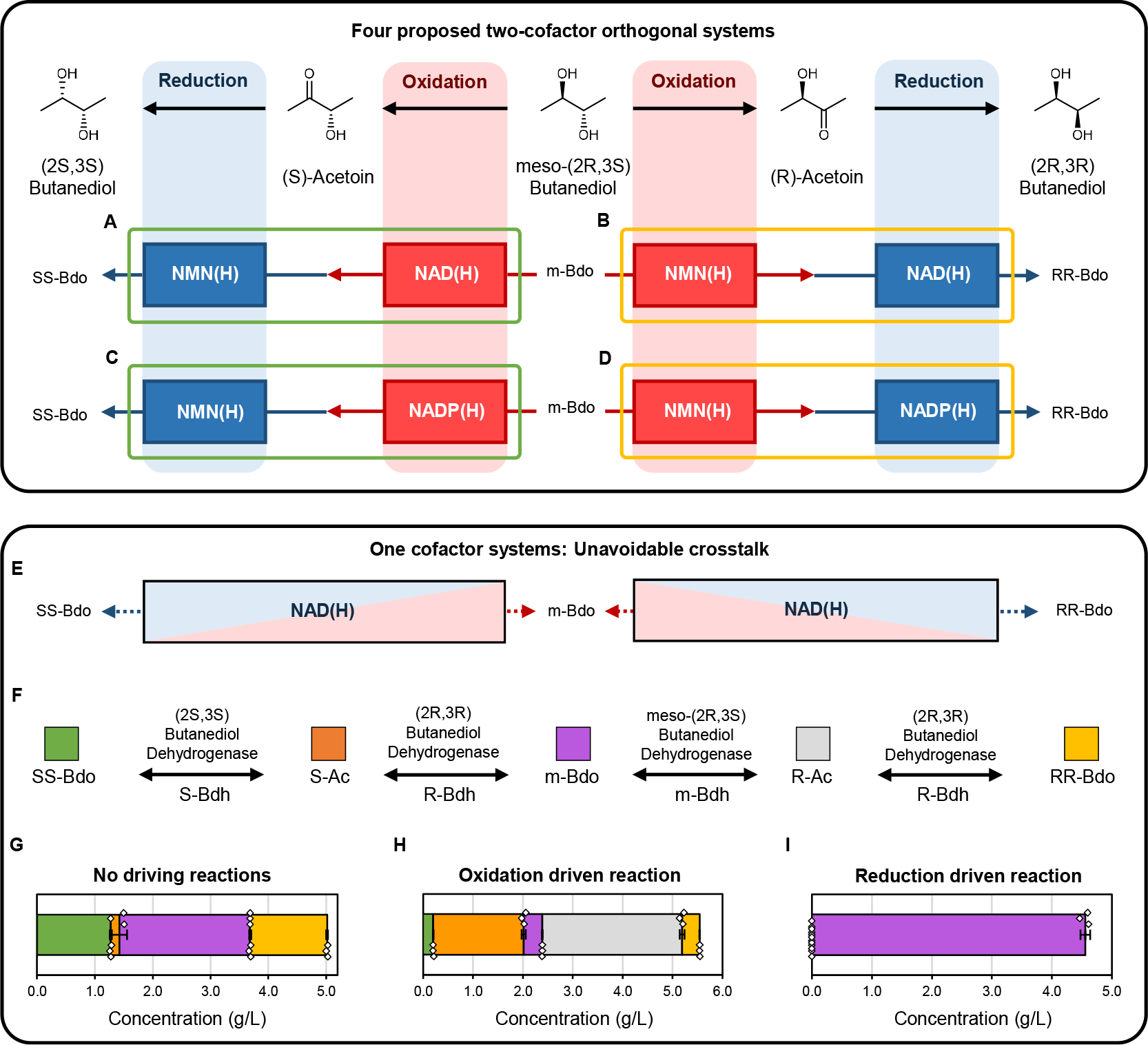
Overview of the four proposed Bdo stereo-upgrading systems for testing two orthogonal cofactor’s crosstalk. **(A, C)** Stereo-upgrading systems of m-Bdo to SS-Bdo where NAD(H) or NADP(H) drive the oxidation step and NMN(H) drives the reduction step. **(B, D)** Stereo-upgrading systems of m-Bdo to RR-Bdo where NMN(H) drives the oxidation step and NAD(H) or NADP(H) drives the reduction step. **(E)** Illustration of when crosstalk between the oxidation and reduction steps occurs, reactions are reversible and incomplete. The four systems above cannot be achieved with a single cofactor or multiple rapidly cross talking cofactors. Red and blue signify oxidation and reduction respectively. The two-color cofactor pool boxes represent rapid crosstalk and fully reversible reaction steps. **(F)** The typical stereospecificity and chiral substrate preference of Bdhs from different enzyme classes used. **(G)** Single cofactor reaction of the Bdhs with no cofactor recycling reactions driving each step. **(H)** Single cofactor reaction with *L. brevis* Nox WT recycling NADH to NAD^+^ to drive the oxidation step. **(I)** Single cofactor reaction with *B. subtilis* Gdh WT recycling NAD^+^ to NADH to drive the reduction step. The reaction does not proceed past the first oxidation step. The substrate for all experiments is 5 g/L m-Bdo and conversion is measured after 48 h at 30 °C. Bars represent the mean of three independent replicates with error bars of one standard deviation. White diamonds indicate the values of individual replicates.

The challenge in this process is that it requires two contradicting steps to both go to completion: First, an oxidation step to destroy a specific chiral center to yield acetoin (Ac); Second, a reduction step to install a new chiral center (Four proposed designs of this process shwn in Fig. 1A, B, C, D, discussed in detail below). If these two steps utilize the same cofactor, for example NAD(H), then the NADH/NAD^+^ ratio would need to be kept low for the first step, and high for the second step. This is not feasible if both steps are occurring simultaneously in the same space (Fig. 1E). When NAD(H) is incubated with 5 g/L m-Bdo and multiple Bdhs with different stereospecificities (different enzyme classes depicted in Fig. 1F), in the absence of any cofactor recycling reactions to drive either redox step to completion, the result is a nearly equal distribution of m-Bdo, SS-Bdo, and RR-Bdo (Fig. 1G). This is because both oxidation and reduction steps suffer from low driving force and operate in a fully reversible fashion. When an oxidation driving force is introduced by coupling to a water-forming NADH oxidase (*L. brevis* Nox WT), the same setup only completes the first oxidation step on m-Bdo, yielding a mixture of S-Ac and R-Ac (Fig. 1H). On the other hand, when a reduction driving force is introduced by NAD^+^-reducing glucose dehydrogenase (*Bacillus subtilis* Gdh WT), the reaction does not proceed past the first step and m-Bdo is untransformed (Fig. 1I).

Broadly, current solutions to this problem in biotechnology are two-fold: First, compartmentalizing the two steps into separate cells with inverse NADH/NAD^+^ ratios^33–35^. This approach requires the intermediate to build up to a significant level and readily diffuse across the cell membranes, which limits the scope of conversion. For example, complex and low concentration intermediates in natural product biosynthetic pathways may not meet these requirements. In vitro, compartmentalization is also difficult to achieve. Second, assigning NADP(H) as the reducing cofactor, and NAD(H) as the oxidizing cofactor. However, although many natural or engineered enzymes can have a preference between NAD(H) and NAD(P)H, most are promiscuous to both cofactors to a substantial degree^36^, due to the intrinsic high similarity between the two cofactors. Therefore, finding or engineering a strongly NAD(H)-specific enzyme for every oxidizing step, and a NADP(H)-specific one for every reducing step, is not always feasible especially as the number of pathway steps increases. Furthermore, the reduction potential of NAD(H) and NADP(H) can become connected in vivo depending on the cell’s metabolic state^37–40^, and NADP(H) is too expensive to use at large scale in vitro^18,37,41–43^.

This model system illustrates that when limited to a single cofactor, metabolic pathways are unable to complete the two contradicting steps no matter how driving force is applied. When two or more cofactors are used with crosstalk among them (e.g. NAD(H) and NADP(H)), the outcome will resemble the single cofactor system to a varying degree, depending on the rate of crosstalk. This justifies a fundamental design principle in metabolism; orthogonality permits thermodynamically incompatible reactions to occur simultaneously.

In our design, we aim to use NMN(H) flexibly as either the reducing or oxidizing cofactor and demonstrate that it can be paired with either NAD(H) or NADP(H) as the opposing cofactor, without crosstalk in the same space. This will result in a total of four combinations in cofactor utilization (Figure 1A, B, C, D). Here, we chose to engineer the enzymes that specifically destroy/install the S-chiral center to use NMN(H) (Fig. 1F). If NMN(H) is an orthogonal redox driving force that is insulated from both NAD(H) and NAD(P)H, two of the four cofactor combinations will yield pure SS-Bdo (Figure 1A, C), and the other two pure RR-Bdo (Figure 1B, D). To demonstrate the versatility of our method compared to the two existing approaches mentioned above, we test both in vitro and in vivo in the same *E. coli* cells, without compartmentalization.

### Development of an Orthogonal NMNH Oxidase

For NMN(H) to drive either oxidation or reduction on demand, our design requires two orthogonal and complimentary NMN(H) cycling enzymes. To maintain a high NMNH/NMN^+^ ratio, we will use our previously engineered NMN^+^-orthogonal glucose dehydrogenase (Gdh Ortho)^8^. To maintain a low NMNH/NMN^+^ ratio, we sought to engineer an orthogonal NMNH-oxidase (Nox Ortho) based on the water-forming NADH-oxidase. Water-forming oxidases (Nox) catalyze the reaction of reduced cofactor with oxygen and provide a byproduct-free and high driving force reaction to rapidly recycle NAD(P)H to NAD(P)^+^ in reported bioprocesses^43–46^. In our recent work, the Nox from *L. pentosus* (*Lp* Nox) has been engineered to accept NMNH and other noncanonical cofactors^26^. However, *Lp* Nox still retains activity toward NADH and NADPH. As discussed above, cofactor promiscuity is undesirable because it breaks the insulation between separate driving forces.

Here, we started with a new Nox scaffold from *L. lactis* due to its high affinity for oxygen, broad operational pH and temperature ranges, and robust applications in vitro and in vivo^47–49^. To enable NMNH activity, we migrated a similar rational design from *Lp* Nox to *Ll* Nox, namely the mutation I159T, which resulted in a ∼20-fold improved activity compared to wild type (WT) with NMNH (Fig. S1A). The potential mechanism of this improvement will be discussed below.

Based on *Ll* Nox I159T, we next sought to further improve NMNH activity by using a high-throughput, growth-based selection platform^26^ (Fig. 2A). This is based on an engineered *E. coli* strain MX 502^26^ (Table S1) that metabolizes glucose exclusively via Gdh Ortho which relies on a NMNH oxidizing enzyme for continuous function. Although this selection is not explicitly designed to yield orthogonal NMNH-oxidases, we hypothesized that orthogonal Nox variants will stand out in the evolution because they will not disturb the NAD(P)H pools essential for cell fitness, and they can turnover NMNH more rapidly without NAD(P)H occupying their active sites as competitive substrates.

**Figure 2.**
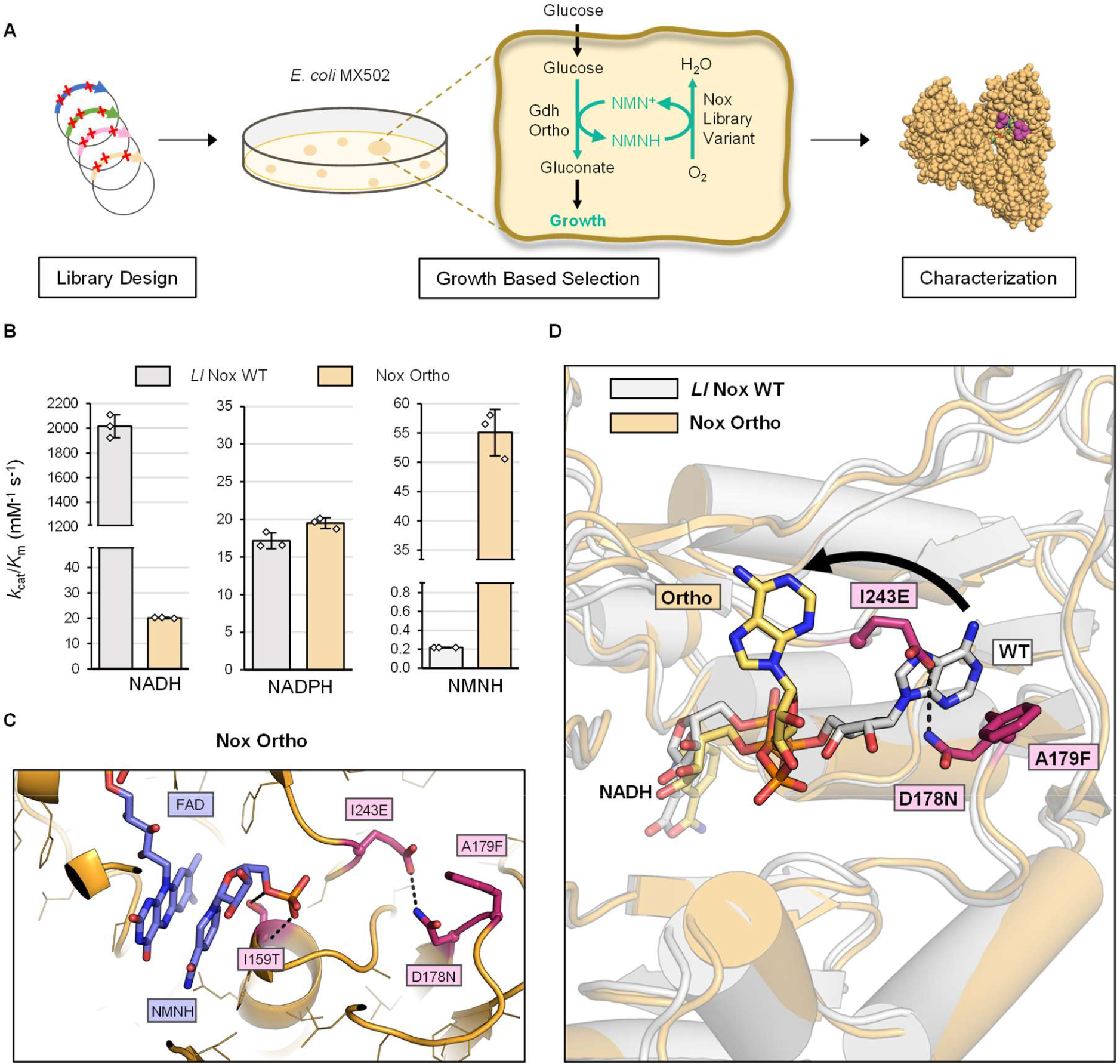
Directed evolution of *Ll* Nox to exclude NADH. **(A)** Schematic of the high-throughput growth-based selection platform workflow for NMNH-utilizing enzymes. The designed site saturated mutagenesis library was introduced to the engineered *E. coli* strain, MX502 where growth depends upon Nox NMNH-oxidase activity. Fast growing variants were characterized by specific activity assay. **(B)** The apparent catalytic efficiencies of *Ll* Nox WT (grey) and Nox Ortho (yellow) towards NADH, NADPH and NMNH. Assay at 37 °C in 50 mM Tris-Cl, pH 7.0 with varying reduced cofactor concentration. **(C)** Model of Nox Ortho binding pose with NMNH and FAD revealed novel hydrogen bond formation. **(D)** *Ll* Nox WT (grey) and Nox Ortho (yellow) with NADH bound revealed a conformational change in NADH binding pose. Mutations on Nox Ortho exclude NADH from its native binding mode to become exposed to solvent, consistent with the decreased catalytic efficiency for NADH observed. Data are presented as the mean of three replicates (n=3) ± one standard deviation.

During growth selection, three positions D178, A179, I243 were subjected to site-saturated mutagenesis with degenerate codons (NNK). These conserved sites had demonstrable effect on the cofactor preference of *Lp* Nox^26^. The constructed library yielded ∼2 × 10^6^ independent transformants, which was sufficient to cover the theoretical library size, 20^3^ = 8000, by more than ten times over. Selection was performed at 30 °C on agar plates of M9 minimal medium supplemented with 20 g/L D-glucose and 2 mM NMN^+^. Plates were monitored for 10 days, and after subsequent colony isolation, eighteen colonies were chosen for further analysis based on desirable growth phenotypes^26^ compared to control plasmids (Table S1). Specific activity assay was performed on sequenced variants (Table S2, Fig. S1B). The best variant *Ll* Nox I159T-D178N-A179F-I243E (Nox Ortho, Table S1) demonstrated a ∼250-fold increase of the apparent catalytic efficiency, *k*_cat_/*K*_m_, toward NMNH (Fig. 2B). Consistent with our hypothesis, Nox Ortho also had the most severely diminished NADH activity, a ∼100-fold decrease in apparent catalytic efficiency compared to WT (Fig. 2B, Table S3). Catalytic efficiency toward NADPH remained low for both WT and Ortho (Fig. 2B, Table S3).

To understand the mechanisms behind Nox Ortho’s drastically switched cofactor preference, Rosetta docking was performed^50^. The resultant models suggested that I159T mutation stabilizes NMNH binding by introduction of a novel polar interaction with the phosphate of NMNH, which is not seen when *Ll* Nox WT was bound with NMNH (Fig. 2C, Fig. S2). D178N and I243E are predicted to form a new hydrogen bond (Fig. 2C, D). A179F potentially restricts the conformer choices of D178N, making the latter sample the hydrogen-bonding conformation more frequently (Fig. 2C, D). This new hydrogen bond is predicted to dislodge NADH from the binding pocket into the solvent (Fig. 2D). This prediction is supported by the very high *K*_m_ of Nox Ortho for NADH and NADPH (Table S3), as these natural cofactors lose contact with the binding pocket.

Reducing the size of cofactor binding pocket is an established approach to engineering enzymes that can utilize smaller, noncanonical cofactors^25–27,51,52^. In fact, in our previous work engineering *Lp* Nox and phosphite dehydrogenase (Ptdh), bulky and hydrophobic residues were selected for at these conserved sites, which pack against one another to fill the cofactor binding pocket and improve noncanonical cofactor activities^26,27^. Those hydrophobic packing interactions, in principle should introduce the same steric hindrance as the hydrogen bond discovered here. Yet, those hydrophobic packing mutations did not effectively block NAD(P)/H binding like the hydrogen bond in Nox Ortho. One explanation may be that a hydrogen bond is much stronger than van der Waals contacts, making the steric hindrance more durable to backbone fluctuation and enzyme conformational shifting during catalysis. Alternatively, when hydrophobic mutations fill the cofactor binding pocket instead of polar contacts, residual binding interaction between the hydrophobic surface of the flexible adenylyl moiety and the hydrophobically packed cleft can persist.

### Development of NMN(H)-Specific Bdo Dehydrogenases

With Gdh Ortho and Nox Ortho in hand, we next needed to design Bdo dehydrogenases that can remove or install the S-chiral center using NMN(H) specifically (Fig. 1A, B, C, D).

For a Bdh that uses NMN(H) as the oxidizing cofactor (Fig. 1B, D), we chose the m-Bdo dehydrogenase from *Klebsiella pneumoniae, Kp* m-Bdh, for its (S)-stereospecificity^53^ and high expression level. Following the same design principle derived from Nox Ortho engineering, we created the NMN(H)-specific variant *Kp* m-Bdh Ortho, which is a triple mutant (M189T-Y34Q-A87K). We predict that M189T will establish novel polar contacts with the terminal phosphate of NMN^+^, and Y34Q-A87K will form a hydrogen bond to close the binding pocket and prevent the entry of NAD(P)^+^ (Fig. 3A, Fig. S3). This design was highly successful; apparent catalytic efficiency of *Kp* m-Bdh Ortho for NMN^+^ improved ∼19-fold compared to WT (Fig. 3B, Table S3). Importantly, *Kp* m-Bdh Ortho’s catalytic efficiency for NAD^+^ and NADP^+^ also decreased 1.6 × 10^5^ and 70-fold compared to WT, respectively. Taken together, *Kp* m-Bdh Ortho features a 3.0 × 10^6^ and 1.3 × 10^3^-fold cofactor specificity switch compared to WT from NAD^+^ or NADP^+^ to NMN^+^, respectively, based on catalytic efficiency (Table S3). Bdh and Nox share little homology (∼13% pairwise sequence identity) and are vastly different structurally. Our success in engineering distinct enzyme families suggests that our design rules are remarkably universal. Current efforts in our lab continue to broaden the classes of enzymes that these rules can be applied to.

**Figure 3.**
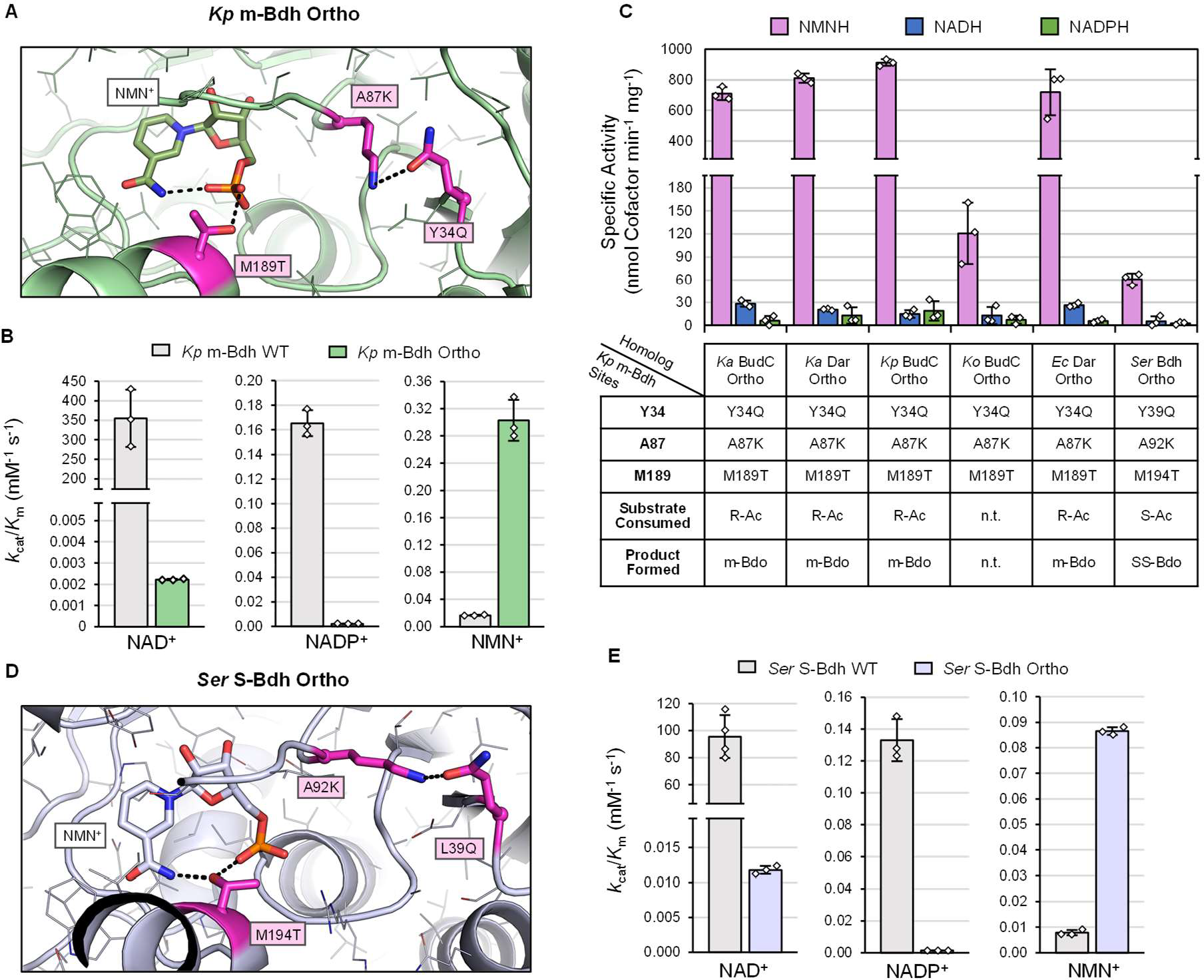
Engineering Bdhs to utilize NMN(H). **(A)** *Kp* m-Bdh Ortho predicted interactions with NMN^+^. **(B)** Apparent catalytic efficiencies of *Kp* m-Bdh WT (grey) and *Kp* m-Bdh Ortho (green) with each cofactor. The substrate was 50 mM m-Bdo with varying cofactor concentration. **(C)** Specific activities of six Bdh homologs based on analogous mutation transfer from *Kp* m-Bdh Ortho (*Kp* m-Bdh sites). Summary of measured chiral substrate and product preference in R/S-Ac feeding experiments (Fig S4). n.t denotes not tested. The substrate was 10 mM R/S-Ac with 0.2 mM reduced cofactor. **(D)** *Ser* S-Bdh Ortho predicted interactions with NMN^+^. **(E)** Apparent catalytic efficiencies of *Ser* S-Bdh WT (grey) and *Ser* S-Bdh Ortho (purple) with each cofactor. Substrate for *Ser* S-Bdh assays was 50 mM SS-Bdo with varying cofactor concentration. Bars are the average of at least three independent replicates with error bars of one standard deviation. Individual replicates are shown as white diamonds. All activity assays of Bdhs conducted at 30 °C in 50 mM Tris-Cl at pH 8.0.

*Kp* m-Bdh Ortho catalyzes the interconversion of m-Bdo and R-Acetoin (R-Ac). For a Bdh that uses NMN(H) as the reducing cofactor (Fig. 1A, C), our stereo-upgrading scheme requires that it accepts S-Acetoin (S-Ac) as substrate to install an additional S-chiral center. To address this design goal, we searched for six other S-installing Bdhs from m-Bdh and S-Bdh enzyme families based on literature or on putative sequence annotation. These Bdhs are: *K. aerogenes* KCTC 2190 BudC, *K. aerogenes* KCTC 2190 Dar, *K. pneumoniae* BudC^54^, *K. oxytoca* KCTC 1686 BudC, *E. cloacae* ssp. *dissolvens* SDM Dar^55^, and *Serratia* sp. AS13^56^ (Table S1, S4, S5). We then rapidly converted all of them into NMN(H)-orthogonal enzymes en masse by mapping the same mutation pattern as applied to *Kp* m-Bdh Ortho onto them (six successful transfers, Fig. 3C). This streamlined process again underscores the facile translatability of our design rules. We determined substrate preference and stereospecificity of all variants by purified protein cycling reactions with NMN^+^ (Fig. S4A). All variants tested exhibit (S)-installing stereospecificity (Fig. 3C, Fig. S4B, C), while only *Ser* Bdh Ortho (L39Q-A92K-M194T) showed the desired substrate preference converting S-Ac into SS-Bdo. Therefore, we renamed this variant *Ser* S-Bdh Ortho.

Rosetta modeling of *Ser* S-Bdh Ortho again supports the predicted mechanism that the hydrogen bond formed by A92K and L39Q, while not affecting NMN^+^ binding (Fig. 3D), pushes NAD^+^ into a solvent-exposed, nonproductive binding pose (Fig. S5). Furthermore, M194T anchors NMN^+^ by interacting with its phosphate (Fig. 3D). Corroborating the computational predictions, *Ser* S-Bdh Ortho (L39Q-A92K-M194T) features an 8.7 × 10^4^ and 1.0 × 10^3^-fold cofactor specificity switch compared to WT from NAD^+^ and NADP^+^ to NMN^+^, respectively, based on apparent catalytic efficiency (Fig. 3E, Table S3).

Taken together, we obtained all necessary parts to assemble the four Bdo stereo-upgrading systems (Fig. 1A, B, C, D): Gdh Ortho to generate NMNH reducing power, Nox Ortho to generate NMN^+^ oxidizing power, *Kp* m-Bdh Ortho and *Ser* S*-*Bdh Ortho to harness these orthogonal driving forces to manipulate (S)-chiral centers.

### Orthogonal Redox Driving Forces Enable Bdo Stereo-Upgrading In Vitro

Using the NMN(H) specific enzymes we developed above and previously^8^, we tested all four designs systematically (Fig. 1A, B, C, D) using purified proteins in vitro (Fig. 4). Gdh WT was used to reduce NAD(P)^+^ and Gdh Ortho was used to reduce NMN^+^. *L. brevis* Nox (*Lb* Nox)^57^, Tp Nox (an *Lb* Nox mutant, Table S1)^39^, and Nox Ortho were used to oxidize NADH, NADPH, and NMNH, respectively. Bdhs of appropriate cofactor, substrate, and chiral specificity were chosen accordingly (additional information in Table S4, S5). Natural cofactors, NAD(P)^+^, were supplemented at 2 mM or 0.2 mM (as noted in Methods), and NMN^+^ at 2 mM. All four systems behaved as intended and demonstrated NMN(H)’s distinct reduction potential compared to NAD(H) and NADP(H) in either oxidizing and reducing directions based on redox ratio (data discussed presented below).

**Figure 4.**
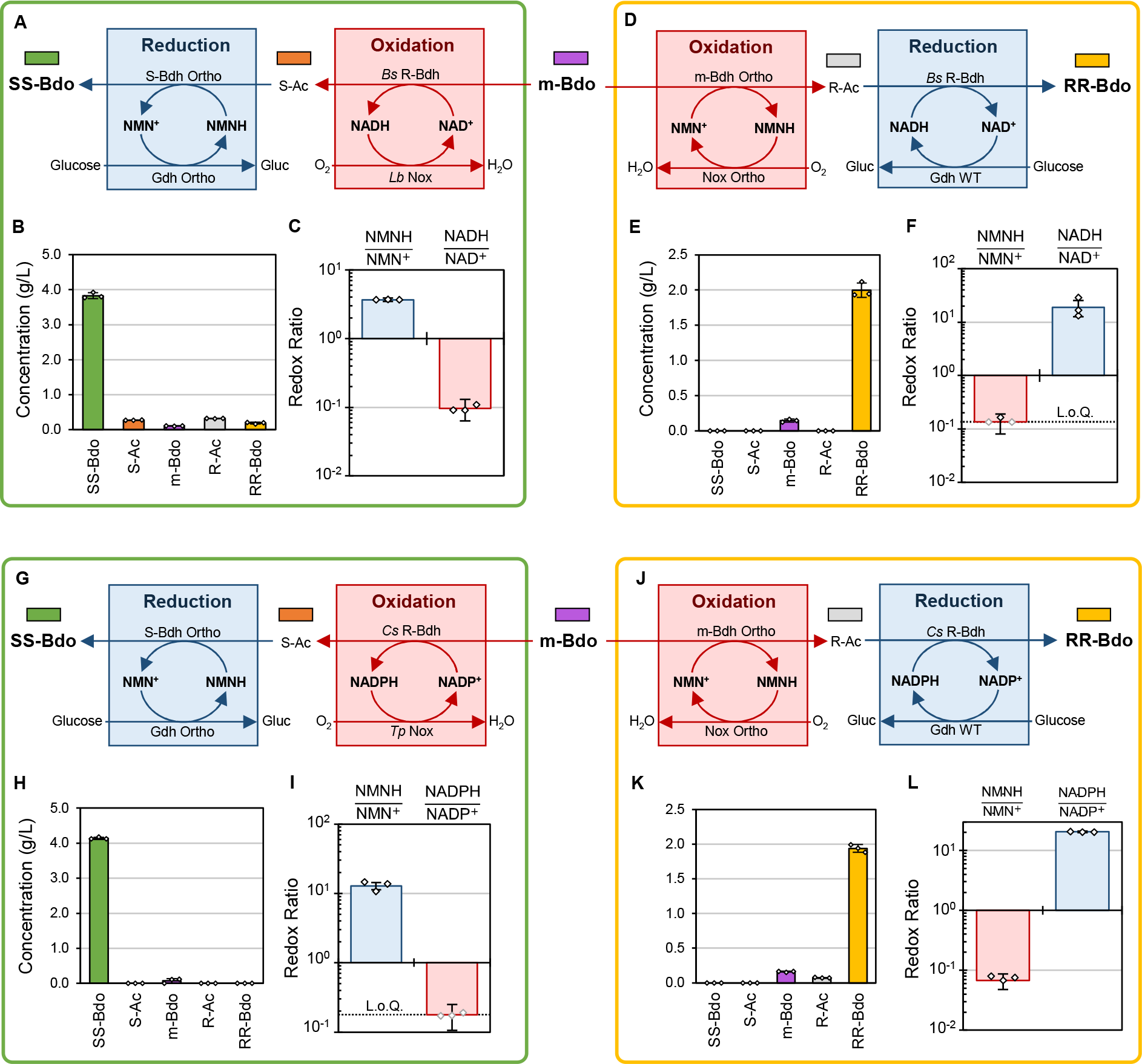
Orthogonal redox driving forces enable Bdo stereo-upgrading in four cell-free systems. **(A-C)** m-Bdo to SS-Bdo system that pairs an NAD(H) driven oxidation step and NMN(H) driven reduction step. **(D-F)** m-Bdo to RR-Bdo system that pairs an NMN(H) driven oxidation step and NAD(H) driven reduction step. **(G-I)** m-Bdo to SS-Bdo system that pairs an NADP(H) driven oxidation step and NMN(H) driven reduction step. **(J-L)** m-Bdo to RR-Bdo system that pairs an NMN(H) driven oxidation step and NADP(H) driven reduction step. **(A, D, G, J)** Reaction pathway maps. **(B, E, H, K)** Concentration of Bdo and Ac isomers. **(C, F, I, L)** Concentration ratio of each redox cofactors’ cognate reduced and oxidized species on a log_10_ scale. Bars represent the average of three independent replicates with error bars calculated by propagation of error from the standard deviation of oxidized and total cofactor concentration measurements. White diamonds with black outline represent the values of individual replicates. Grey outlined diamonds represent replicates whose reduced cofactor concentration is below limit of quantification (L.o.Q., see Methods). Ortho represents the cofactor engineered variants. Gluc, gluconic acid. All reactions incubated shaking for 24 h or 72 h at 30 °C for SS-Bdo or RR-Bdo production, respectively. Samples for redox ratio measurement were taken from the same reactions at the times listed in Methods.

In the system that included *Lb* Nox, *Bacillus subtilis* R-Bdh (*Bs* R-Bdh), Gdh Ortho, and *Ser* S-Bdh Ortho (Fig. 4A, 1A), 5 g/L of m-Bdo was converted to ∼3.8 g/L SS-Bdo (77% conversion) (Fig. 4B). Here, NMN(H) is tasked to be the reducing power and we observe a high NMNH/NMN^+^ ratio at 3.7 ± 0.2 (Fig. 4C), while NADH/NAD^+^ was maintained at a low ratio, 0.10 ± 0.03. On the other hand, in the system that included Nox Ortho, *Kp* m-Bdh Ortho, Gdh WT, and *Bs* R-Bdh (Fig. 4D, 1B), 2 g/L of m-Bdo was converted to ∼1.9 g/L RR-Bdo (93% conversion) (Fig. 4E). Here, NMN(H) played the role of the oxidizing power with a low NMNH/NMN^+^ ratio of ≤ 0.14 ± 0.05, based on NMNH levels near limit of quantification for this system (L.o.Q., 200 µM, see Methods), versus NADH/NAD^+^ at 19 ± 6 (Fig. 4F). Both systems achieved high purity in producing the desired Bdo isomers. The stark contrast between NMN(H) and NAD(H) redox ratios supports that these two cofactors are orthogonal to each other.

The further the distance between the two cofactors’ redox ratios, the more strongly they can deliver two opposing, insulated driving forces; this insulation is ensured by all enzymes’ strict cofactor specificities. Indeed, when comparing the SS-Bdo producing system (Fig. 4A, B, C, Fig. 1A) to the RR-Bdo producing system (Fig 4D, E, F, Fig. 1B), we observed that the closer NMN(H) versus NAD(H) redox ratios in the SS-Bdo producing system resulted in *Bs* R-Bdh and *Ser* S-Bdh Ortho retaining small residual ability to catalyze the reverse, unintended reactions, as evident by the slight buildup of side products RR-Bdo, R-Ac, and S-Ac. This could be due to the small side activity of *Ser* S-Bdh Ortho toward NAD^+^ (Fig 3E, Table S3), which penetrates the insulation and permits some slow leakage by crosstalk between reactions. This was not observed in the system based upon *Kp* m-Bdh Ortho paired with NAD(H) (Fig 4D, E, F, Fig. 1B), which has superior specificity for NMN^+^ (Fig. 3B, Table S3). This leakage phenomenon due to crosstalk between reactions by shared cofactor usage exists and is shown to be prevalent and profound between the natural cofactors, NAD(H) and NADP(H), in Nature, because of the presence of numerous promiscuous enzymes^36^ and dynamic use of direct and indirect modes of cofactor exchange^6,37,39,40,58^. The high structural deviancy of noncanonical cofactors promises to mitigate this.

After showcasing our Bdo stereo-upgrading system’s capacity to orchestrate orthogonal redox driving forces from complex mixtures utilizing both NAD(H) and NMN(H). We next sought to demonstrate that these principles are translatable to systems based on the other natural redox driving force, NADP(H), and thereby compatible with direct implementation in vivo. The system using NADP(H) paired with NMN(H) was also prepared. Previous literature described NADPH-dependent R-Bdhs^59–61^, but we found their activity in m-Bdo oxidation to be limited. Therefore, we bioprospected an NADP^+^-active R-Bdh from *Clostridium saccharoperbutylacetonicum* (*Cs* R-Bdh) based on sequence homology to the readily reversible *Bs* R-Bdh (Fig. S6A-C). We subsequently confirmed our R-Bdh’s specific activity with NADP^+^ and stereospecific oxidation to destroy the R-chiral center of m-Bdo (Fig. S6D-G).

The SS-Bdo producing system contained Tp Nox, *Cs* R-Bdh, Gdh Ortho, and *Ser* S-Bdh Ortho (Fig. 4G), and produced ∼4.1 g/L SS-Bdo from 5g/L m-Bdo (83 % conversion) (Fig. 4H). The redox ratios suggest NMN(H) can be held far from equilibrium with NADP(H), with NMNH serving as a strong reducing power (NMNH/NMN^+^ = 13 ± 2) (Fig. 4I), while the NADP(H) pool stays oxidized at < 0.18 ± 0.07 calculated based on our NADPH limit of quantification in this system (260 µM, see Methods), and NADP^+^ levels consistent with the total concentration provided, Fig. 4I). Unlike the previous SS-Bdo producing system based on combination with NAD(H) (Fig. 4A), the NAD(P) system (Fig 4G) does not suffer byproduct formation; consistent with the *Ser* S-Bdh Ortho’s nearly perfect orthogonality toward NADP^+^ (Fig 3B, Table S3). The complementary stereo-upgrading system contained Nox Ortho, *Kp* m-Bdh Ortho, Gdh WT, and *Cs* R-Bdh (Fig. 4J) and produced ∼1.9 g/L of RR-Bdo from 2 g/L m-Bdo (89 % conversion) (Fig. 4K). Here, with NMN(H) as the oxidizing power, the NMNH/NMN^+^ ratio was expectedly low (0.07 ± 0.02), while the reducing power in this system, NADP(H), was high (NADPH/NADP^+^ = 20. ± 0.6) (Fig. 4L). These results demonstrate ideal orthogonality between NADP(H) and NMN(H) for the systems based on *Kp* m-Bdh Ortho.

Altogether, *Kp* m-Bdh Ortho, *Ser* S-Bdh Ortho, and Nox Ortho which share a unifying enzyme design principle for cofactor specificity, combine to form four different stereo-upgrading systems capable of remarkedly pure preparations of chiral Bdo.

### Orthogonal Redox Driving Forces Enable Bdo Stereo-Upgrading in *E. coli* Whole Cells

To convert m-Bdo to SS-Bdo we start with a previously reported *E. coli* strain^8^, MX102 R^0^ (Table S1). This cell is unable to catabolize glucose, our sacrificial electron donor, and has decreased ability to degrade NMN^+8,62^ (Fig. 5A). We first used NAD(H) as the oxidant and NMN(H) the reductant (Fig. 5B). *Lb* Nox, *Bs* R-Bdh, Gdh Ortho, *Ser* S-Bdh Ortho, and *Zymomonas mobilis* (*Zm* Glf) encoding a glucose transport facilitator are coexpressed from plasmids (Fig. 5B, pDA129, pDA131, and pSM10 in Table S1) in *E. coli* resting cells. When cells are incubated with 200 mM glucose, 5 g/L m-Bdo, and 10 mM NMN^+^ supplemented, we were able to produce 3.8 g/L SS-Bdo (Fig. 5C). The product purity of cells supplemented with 10 mM NMN^+^ also dramatically increased relative to cells without NMN^+^ supplementation, with the former reaching 81% pure SS-Bdo (Fig. 5D). When no NMN^+^ was supplemented, substantial amount of S-Ac was observed. This is indicative of undesirable reversible reactions caused by insufficient driving forces, as discussed above.

**Figure 5.**
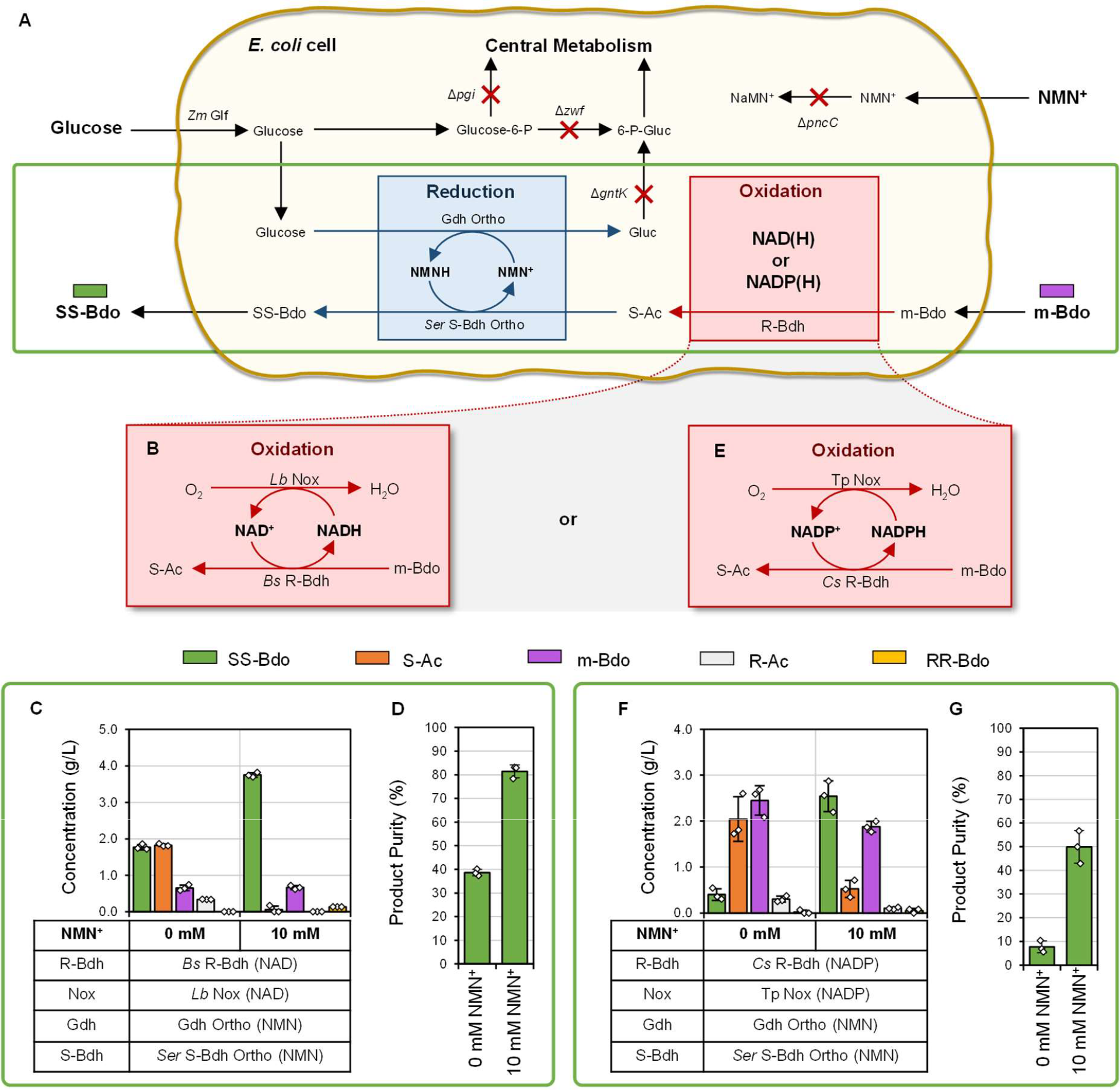
Resting cell stereo-upgrading of m-Bdo to SS-Bdo in *E. coli*. **(A)** Gene deletions and reaction pathways in the whole cell chassis, strain MX 102 R^0^. **(B-D)** m-Bdo to SS-Bdo system using NAD(H) driven oxidation step paired with an NMN(H) driven reduction step in vivo. **(B)** Reaction pathway details of the NAD(H) driven oxidation step. **(C)** Concentration of Bdo and Ac stereoisomers when 0 mM or 10 mM NMN^+^ is supplemented to the media, sampled after 24 h at 30 °C. **(D)** Product purity of SS-Bdo in different NMN^+^ supplementation conditions. **(E-G)** m-Bdo to SS-Bdo system using NADP(H) driven oxidation step paired with an NMN(H) driven reduction step in vivo. **(E)** Reaction pathway details of the NADP(H) driven oxidation step. **(F)** Concentration of Bdo and Ac stereoisomers when 0 mM or 10 mM NMN^+^ is supplemented to the media, sampled after 152 h at 18 °C. **(G)** Product purity of SS-Bdo in different NMN^+^ supplementation conditions. Bars represent an average of three biological replicates with error bars of one standard deviation. White diamonds represent the values of individual replicates. Product purity calculated as the percentage of SS-Bdo in the total amount of products formed (R-Ac, S-Ac, RR-Bdo, SS-Bdo, m-Bdo). Ortho represents the cofactor engineered variants. Gluc, gluconic acid; 6-P-Gluc, 6-phosphogluconate; Glucose-6P, glucose-6-phosphate; NaMN, nicotinic acid mononucleotide.

The system can also function with NADP(H) as the oxidant, when *Lb* Nox is replaced with Tp Nox, and *Bs* R-Bdh is replaced with *Cs* R-Bdh (Fig. 5E, pDA129, pDA162, and pSM10 in Table S1). The resting cells produced a final titer of 2.5 g/L SS-Bdo when 10 mM NMN^+^ is supplied (Fig 5F). Without NMN^+^ supplementation, very little SS-Bdo is produced (0.40 g/L) even though a high concentration of S-Ac is formed (2.0 g/L), suggesting a lack of reducing power when NMN^+^ is not supplied. Again, the product purity in the NMN^+^ supplied cells was greatly improved, from 7.7% pure with 0 mM NMN^+^ to 50.% pure SS-Bdo with 10 mM NMN^+^ (Fig. 5G). The primary impurity in this system is m-Bdo, the starting material, which points to the first, oxidative step catalyzed by the NADP(H)-active *Cs* Bdh and driven by NADPH-specific Tp Nox as the major bottleneck. The equivalent cell-free system readily catalyzes the oxidation with NADP^+^ (Fig. 4G, H, I). Yet, even in resting *E. coli* cells, the redox ratio of NADP(H) is challenging to invert toward efficient oxidation. The small total NADP(H) pool size could also impact the efficiency of this step; basal NADP(H) concentrations are on the order of 0.1 mM^63^ whereas 2 mM NADP^+^ was provided in the cell-free system (Fig. 4G, H, I).

Orthogonal metabolic systems can vastly simplify the optimization efforts required to achieve whole cell biotransformation, because the engineering efforts can be primarily focused on the heterologous pathway itself. Here, our orthogonal system achieves high product purity with minimal alterations to the host’s native metabolism. This work establishes that NMN(H) reduction potential can be firmly enforced in vivo without the interference from the NAD(H) and NADP(H) redox state, which will deliver persistent driving force in the ever-changing cellular environment.

## Conclusion

This work uncovers an enzyme design principle, polar contacts that bridge across the cofactor binding pocket as a reliable strategy to engineer NMN(H)-orthogonal biocatalysts. Parallel implementation of this design principle across diverse enzymes enables the construction of orthogonal metabolic systems. Here, we recreate one of the most important design principles in cellular metabolism, chemical compartmentalization by specific coenzymes^39,64^ and quantify actual reduction potentials by the redox ratios of cofactors in our two-dimensional redox systems. The methodology presented here on a synthetic redox metabolism is true to the seminal description of orthogonal metabolic organization present in Nature^64^ .

Additionally, we show that the reduction potential of NMNH/NMN^+^ is disconnected to that of NADH/NAD^+^ and NADPH/NADP^+^, in both cell-free preparations and *E. coli* whole-cells. Partitioning and balancing redox cofactors is a prevalent need in metabolic engineering. Unlike NAD(P)^+^, noncanonical cofactors are not tied to natural metabolism. Given that their recycling enzymes, such as Gdh Ortho and Nox Ortho, can be expressed in the desired cellular compartment, with tunable level, and at a controllable timing; noncanonical redox systems can support precise spatiotemporal control. Therefore, they may be deployed to solve the prevalent problem of balancing redox in metabolic engineering with a much wider range of freedom. This work presents the full infrastructure of using NMN^+^ as an orthogonal redox cofactor with its designated electron source and electron sink. We demonstrate this concept using NMN(H) to direct unwavering redox driving forces using Bdo stereo-upgrading as a proof-of-concept. Looking forward, access to intricate control of reduction potential in individual metabolic processes, in separate cellular or organ compartments, at the correct time, and without interference with delicate global redox homeostasis is essential^57,58,65–68^; precise maintenance of the delicate redox patterns may promote health and prevent diseases^58^.

## Methods

### Media and Growth Conditions

Culture for cloning was routinely performed with *E. coli* XL-1 Blue strain (Stratagene) and protein expression was generally carried out in *E. coli* BL21 (DE3) strain (Invitrogen). Unless otherwise noted all *E. coli* cultures were grown in 2x YT medium prepared from ready-to-use media granules at 31 g/L (Fisher Bioreagents 2x YT Broth granulated media: 16 g/L casein peptone, 10 g/L yeast extract, 5 g/L NaCl). When appropriate, selective media was prepared at working concentrations of ampicillin (100 mg/L), spectinomycin (50 mg/L), kanamycin (50 mg/L), or chloramphenicol (20 mg/L from an ethanol solvated stock). Generally, strains were cultured at 37 °C with 250 rpm agitation on a 1-inch diameter orbit, and induction was initiated with final concentrations of 0.1% L-arabinose for strains containing the *P*_*BAD*_ promoter and 0.5 mM isopropyl ß-D-1-thiogalactopyranoside (IPTG) for strains containing the *P*_*LlacO1*_ promoter, unless otherwise noted. Buffer compositions are detailed in the Supporting Information.

### Plasmid Construction

Standard PCR reactions were performed using the PrimeSTAR Max DNA Polymerase (TaKaRa) or KOD One PCR Master Mix -Blue-(Toyobo). Splicing by overlap extension PCR was performed with the KOD Xtreme Hot Start DNA Polymerase (Novagen). Generally, plasmids were constructed by PCR amplification of the target gene, with appropriate ∼20-30 bp overlapping regions for downstream Gibson assembly^69^ included in the primers. Constructs were circularized with vector backbone by Gibson assembly and introduced to XL-1 Blue Mix and Go chemical competent cells (Zymo Research), miniprepped (Qiagen), and sequenced (Laragen). pDA63 was constructed by amplification of the *bdhA* gene from a *Bacillus subtilis* ATCC 6051 genome template. Plasmids containing previously reported genes were subcloned from plasmids^70^ (Table S1).

Reported plasmid constructs from synthetic genes such as *L. lactis nox* were ordered from Integrated DNA Technologies (IDTDNA) as gBlocks gene fragments with codon optimization performed by the IDTDNA codon optimization webtool for *E. coli* K12. All bioprospected genes were prepared as synthetic DNA. DNA fragments were designed with the appropriate ∼30 bp Gibson overlapping sequences on each end for direct circularization into vector backbones by Gibson assembly. Plasmids encoding mutants were built through site-directed mutagenesis as previously described^8^ or encoded in the synthetic DNA design.

### Strain Construction

Strain MX102 R^0^ was constructed by removal of the kanamycin resistance marker using the pCP20 plasmid (Table S1) from the previously reported strain^8^.

### Protein Expression and Purification

Expression of recombinant protein was performed by introduction of the target plasmid into BL21(DE3) chemically competent cells (Table S1). A single colony was inoculated into 4 mL 2x YT-200 mg/L ampicillin and grown for ∼14-16 h at 30 °C. 2x YT-200 mg/L ampicillin media was inoculated to 0.07 OD_600_ and incubated while shaking for approximately 1 h and 45 min at 37 ºC. The cultures were removed from the shaker and placed at room temperature for 30 min without shaking, induced with 0.5 mM IPTG, then incubated while shaking at 30 ºC, 24 h. Cells were harvested by centrifugation at 2500 *g*, 4 ºC, 30 min. Supernatant was decanted, and cell pellets stored at -80 ºC until purification. Cell lysis was performed mechanically, and purification was performed following a modified version of the Zymo His-Protein Miniprep detailed in the Supporting Information. Protein quantification was achieved by Bradford assay relative to a standard curve of bovine serum albumin. Purified protein was stored with 20% glycerol at -80 ºC.

### Preparation and Purification of NMNH

NMNH was prepared enzymatically and purified as described previously^26^.

### Specific Activity Assay

Specific activity assays performed to characterize *Kp* m-Bdh homologs were done in the following conditions: 50 mM Tris-Cl at pH 8.0, R/S-Ac at 10 mM, and 0.2 mM reduced cofactor, 30 °C. 100 µL reactions were prepared by addition of master mix to 10 µL of each enzyme. Enzyme stored in His-Elution Buffer with 20 % glycerol (from a 50 % glycerol stock). Enzyme dilutions were carried out with ice cold His-Elution Buffer. Initial reaction rates were determined from the slope of the first 60-100 s of reaction. Each sample was assayed in at least three replicates, unless otherwise noted and the reaction rates are recorded as the difference between substrate and no substrate control. Production or consumption of reduced cofactor was detected with a SpectraMax M3 spectrophotometer at 340 nm, where the extinction coefficient used for calculation of NAD(P)H reaction rate was 6.22 mM^-1^ cm^-1^ or 4.89 mM^-1^ cm^-1^ for NMNH.

Specific activity assays performed to characterize the *Bs* R-Bdh homologs were carried out in the following conditions: 50 mM glycine-NaOH pH 10.0, 10 mM m-Bdo, 4 mM oxidized cofactor. Specific activity assays of Nox variants were carried out in the following conditions: 50 mM Tris-Cl at pH 7.0 with 0.3 mM reduced cofactor at 37 °C. The reaction rate was calculated as the difference between no cofactor control and with cofactor reaction to account for potential non-specific absorbance decrease at 340 nm in the absence of the addition of reduced cofactor. The dilution buffer for Nox was 50 mM Tris-Cl at pH 7.0. Unless otherwise noted, handling of enzymes, initiation of reaction, and calculation of reaction velocity was performed as described above.

### Apparent Kinetic Parameter Determination

Apparent Michaelis-Menten kinetic parameters for cofactors were determined by nonlinear fit of recorded initial reaction rate data to the Michaelis-Menten equation below – where *v*_*0*_ is initial rate, *E*_T_ is the total enzyme concentration, and *C* is the varying cofactor concentration. For Bdhs the kinetic parameters, *k*_cat_ and *K*_m,_ describe the apparent turnover number and apparent Michaelis constant at 50 mM substrate, respectively.

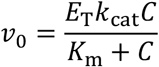

For *Ser* S-Bdh the assay condition was 30 °C, 50 mM Tris-Cl pH 8.0, 50 mM SS-Bdo, and cofactor concentration varied. For *Kp* m-Bdh the same assay condition was performed but with 50 mM m-Bdo. For *Ll* Nox the assay condition was 37 °C, 50 mM Tris-Cl pH 7.0 with variable reduced cofactor concentration.

Under conditions where the enzyme could not be saturated with cofactor (*K*_m_ ≫ *C*) the initial rate data were instead recorded and fit to a linear form of the Michaelis-Menten equation to solve for catalytic efficiency, *k*_cat_/*K*_m_.

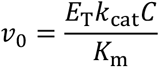

### Gas Chromatography

All gas chromatography (GC) was performed with an Agilent 6850 (Agilent Technologies) coupled to a flame-ionization detector. All gases were purchased from Airgas. The Agilent CP-ChiraSilDex CB (25 m x 0.25 mm ID x 0.25 µm film thickness x 5-inch cage) was used for all separations. A method was developed with the following parameters: The inlet was heated to 250 °C with a pressure of 25.0 psi. Injection volume was 1 µL. The inlet was set to split at a ratio of 20:1 with helium used as the carrier gas. The GC was operated in constant pressure mode at 25.0 psi. The oven program started at 80 °C, hold for 10 min, then 10 °C/min ramp to 110 °C, followed by 20 °C/min ramp to 200 °C with a 2 min hold. The flame ionization detector was set to 275 °C, 40 mL/min H_2_, 350 mL/min air, with a makeup helium flow of 45 ml/min. The elution order observed is R-Ac, S-Ac, SS-Bdo, RR-Bdo, m-Bdo, and then internal standard, 4-oxoisophorone. Samples for GC analysis were processed by a salting-out-extraction with ethyl acetate as detailed in Supporting Information.

To determine the retention times of the Bdo stereoisomers, an analytical standard of each prepared in ethyl acetate was injected and compared to reaction chromatograms. For Ac, both literature retention order on an equivalent column chemistry and a reaction mixture of an enzyme with published (S)-chiral center installing stereospecificity^53^ was used to generate pure (S)-acetoin from diacetyl and record a characteristic retention time. Integrated peak area for each analyte and internal standard was recorded for each sample. A calibration curve was generated for the response ratio (area analyte/area of internal standard) of standards containing known mixtures of all Bdo and Ac stereoisomers with fixed internal standard concentration. Concentrations of each were interpolated from standard curves, calculated by linear fit.

### Purified Protein Cycling Reactions

Cycling reactions consisting of purified proteins were prepared similarly to previous research^8^. Reactions were initiated by addition of a mixture of each reaction’s enzymes to a master mix containing all other components. For determination of enzyme stereospecificity, reaction mixtures contained 100 mM potassium phosphate (KP_i_) pH 7.5, 1 M NaCl, 200 mM D-glucose, 2 mM oxidized cofactor, and 5 g/L racemic acetoin. The final concentration of enzyme in the reaction was 11.7 µM Gdh Ortho and 28.1 µM of Bdh. All molar protein concentrations are reported based on the molecular weight of the monomers. When an enzyme was omitted from a reaction, an equivalent volume of His-Elution Buffer was added in its place. The 350 µL reactions were incubated in 15 mL conical tubes at a 45° angle, 30 °C, 250 rpm for 4 h prior to GC analysis as described above. Analogous reactions for R-Bdh bioprospecting are described in the Supporting Information.

For the single cofactor model systems, 5 g/L m-Bdo, 2 mM NAD^+^, 100 mM KP_i_, 200 mM 3-morpholinopropane-1-sulfonic acid (MOPS), pH 8.0, 10 µM *Ser* S-Bdh WT, 5 µM *Bs* R-Bdh were provided. Cofactor recycling enzymes consisted of 10 µM *Lb* Nox or 10 µM Gdh WT as appropriate. 450 µL reactions were incubated while shaking for 48 h at 30 °C prior to sample analysis by GC.

For the m-Bdo to SS-Bdo reaction cascade using NAD^+^ and NMN^+^ (Fig. 1A, 4A), the 900 µL reactions contained 300 mM Tris-Cl pH 8.0, 1 M NaCl, 200 mM D-glucose, 2 mM oxidized cofactor, and 5 g/L m-Bdo. The protein ratio used was 10 µM *Lb* Nox, 5 µM *Bs* R-Bdh, 20 µM Gdh Ortho, and 60 µM *Ser* S-Bdh Ortho. For the NADP^+^ and NMN^+^ system (Fig. 1C, 4G), the protein ratio used was 20 µM *Tp* Nox, 10 µM *Cs* R-Bdh, 20 µM Gdh Ortho, and 60 µM *Ser* S-Bdh Ortho. Reactions to produce SS-Bdo were sampled after 24 h at 30°C, for GC analysis.

For the m-Bdo to RR-Bdo reaction cascade using NAD^+^ and NMN^+^ the 900 µL reactions contained 300 mM Tris-Cl pH 8.0, 1 M NaCl, 200 mM D-glucose, 2 mM NMN^+^, 0.2 mM NAD^+^ and 2 g/L m-Bdo. Since determination of *K*_m_ for Nox Ortho with NAD(P)H was technically limited, specific activity assay of Nox Ortho with each cofactor prototyped to guide system design (Fig. S7). The concentration of NAD(P)^+^ in these systems was 0.2 mM. Protein ratio was 10 µM Nox Ortho, 60 µM *Kp* S-Bdh Ortho, 5 µM *Bs* Gdh, and 5 µM *Bs* R-Bdh. For the NADP^+^ and NMN^+^ system, NADP^+^ was substituted for NAD^+^ and 5 µM *Cs* R-Bdh was substituted for *Bs* R-Bdh. Reactions to produce RR-Bdo were sampled after 72 h at °C, for GC analysis.

### Standard Selection Protocol of *Ll* NOX Libraries and Control Plasmids into MX502

The strain construction and standard selection were performed as described previously^26^. Plasmid pLS501 (XenA D116E) and pYZ10 (*Ll* Nox WT) served as positive and negative controls, respectively. Plasmid pYZ20 (*Ll* Nox I159T) was also transformed as library template control as a benchmark.

### Selection of *Ll* Nox Library pYZ100

The library culture was obtained by following the standard selection protocol described above and was washed three times in M9 Wash Buffer, was diluted to a final concentration of ∼10^5^ cells/mL in M9 Wash Buffer. 40 µL of this suspension was plated on each of seven M9 Selection Plates with 2 mM NMN^+^. Plates were incubated at 30 °C and monitored periodically for 10 days. After 10 days, six colonies from each plate and colonies from the control plates were streaked onto fresh M9 Selection Plates to isolate variants and validate growth. After 7 days, plates were evaluated for growth and 18 colonies with diverse growth phenotypes were selected for sequencing (Table S2) and NMNH activity characterization. Colonies were cultured in 2x YT liquid media overnight and plasmids were extracted using QIAprep Spin Miniprep kit (Qiagen).

### Reformulation of Commercial Redox Ratio Kits for NMNH/NMN^+^ quantification

The Amplite Colorimetric NADH Assay Kit (AAT Bioquest) with some components from the redox ratio kits were reformulated to prepare an NMNH/NMN^+^ kit. The NAD^+^/NADH ratio and NADP^+^/NADPH ratio quantification was performed according to the kit’s instructions with additional pre-assay sample processing steps to remove enzymes detailed in supporting information. The NMNH/NMN^+^ reformulated kit method is detailed in the supporting information. Briefly, the difference between the NADH assay kit and the redox ratio kits is the addition of a selective reduced cofactor degradation step and cofactor specific enzymatic recycling. Therefore, we prepared the NMNH/NMN^+^ ratio quantification kit by using the selective cofactor degradation solutions from the redox ratio kits with addition of a strictly NMN^+^ specific reducing enzyme, Gdh Ortho^8^, to the final colorimetric assay master mix solution. Data for Fig 4C, F, I, L which correspond to systems in Fig 1A, B, C, D were sampled at 48 h, 72 h, 48 h, and 24 h respectively.

### Limit of Reduced Cofactor Quantification (L.o.Q.)

The commercial and reformulated redox ratio kits have higher variability for the reduced cofactor, which complicates quantification of low reduced cofactor concentrations and low redox ratios. This is due to the indirect measurement of the reduced cofactor concentration, where the reduced cofactor concentration is calculated as the difference between the concentration of a specific oxidized cofactor (e.g. NADP^+^) and its total concentration (e.g. NADP(H)). To ensure we report redox ratios, calculated from reduced cofactor concentrations, that are confidently quantifiable and do not underreport a given redox ratio, we define the L.o.Q. for each reduced cofactor species in each system based on a 95% confidence level for *n* samples in the difference of two measured means, of oxidized and total cofactor concentration, with corresponding measurement variability of one standard deviation, σ_ox_ and σ_total_. Here, the 95% confidence level is calculated using the *t* statistic for *n*=3, *t*_0.95_.

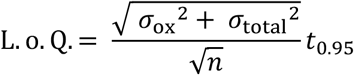

For an individual replicate, *x*, the reported reduced cofactor concentration, and redox ratio are calculated by the following equation, where values calculated from concentrations set at L.o.Q. are indicated with grey outlined diamonds.

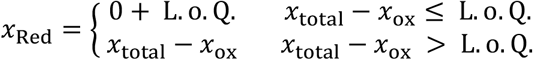

The mean reduced cofactor concentration is also calculated with each replicate’s *x*_Red_ determined as described.

### Resting Cell Biotransformation

All necessary plasmids were inserted into strain MX102 R^0^ by electroporation and plated on 2x YT with 200 mg/L ampicillin, 50 mg/L spectinomycin, and 50 mg/L kanamycin (ASK). Triplicate seed cultures of 4 mL 2x YT-ASK with 5 g/L D-mannitol were prepared from single colonies and grown at 37 °C for 16 h. Cell cultivation cultures were normalized to 0.04 OD_600_ in 50 mL 2x YT-ASK with 10 g/L D-mannitol and 0.5 mM 5-aminolevunic acid. The cells were incubated shaking for 4 h at 30 °C and then induced with 0.5 mM IPTG and 0.1% (w/v) L-arabinose. After induction, cells were incubated for 12 h while shaking at 30 °C before harvest. At harvest the OD_600_ of each culture was determined and the volume of cells necessary to prepare a final suspension of 100 OD_600_ in 3 mL final volume was transferred to 50 mL conical tubes and pelleted for 10 min at room temperature, 2500 *g*. The pelleted cells were washed by gentle resuspension and centrifugation twice in 40 mL 100 mM KP_i_ pH 7.5 at room temperature, then washed one more time in 10 mL of the same and transferred to 15 mL conical tubes. The conical tubes were pelleted a final time and resuspended to a total volume of 3 mL.

The resting cell reactions were initiated by the addition of 600 µL washed cell suspension to 2.4 mL concentrated reaction master mix in 15 mL conical tubes. Reaction mixture was composed of 200 mM D-glucose, 0 or 10 mM NMN^+^, 5 g/L m-Bdo, 0.5 mM IPTG, 100 mM KP_i_ dibasic, 200 mM MOPS, ASK, and 0.1% (w/v) L-arabinose. Resting cell reactions were then incubated horizontally at 30 °C, 250 rpm, for 48 h. Resting cell reactions with Tp Nox expressed were instead incubated at 18 °C for 152 h.

### Rosetta Molecular Simulations

The *Ll* Nox, *Kp* m-Bdh, and *Ser* S-Bdh structures were generated by Alphafold^71^ and subsequently relaxed using a backbone constrained FastRelax procedure^72^. The NMN^+^, NAD^+^, and NADP^+^ conformer library used was previously established^28^. Initial coordinates for NMN^+^ were obtained by aligning the *Ll* Nox structure to the homologous structure PDB: 2BC0^73^. Initial coordinates forNMN^+^ in *Kp* m-Bdh and *Ser* S-Bdh were obtained by alignment to the homologous structures PDB: 1GEG^54^ and PDB: 3A28^74^, respectively.

Detailed method for *Ll* Nox is given below. Bdhs were simulated in the same manner with the exclusion of FAD^+^. For each simulation, one of the cofactors and FAD^+^ were placed into the active site, and the enzyme-ligand complex was optimized. The EnzRepackMinimize protocol was applied to predict the WT structure bound with NMN^+^ and amino acid substitutions that improve binding affinity for NMN^+75^. This process involved Monte Carlo evaluation, sampling alternative rotamers, side chain substitutions, and backbone minimization to examine different binding pocket geometries. Restraints were imposed to maintain the cofactors in a catalytically competent geometry for hydride transfer and interactions with FAD^+^. After 2000 simulations for each batch, the top 20 outputs based on constraint score, protein-ligand interface energy score, and total system energy score were visually examined. For ligand docking simulation with mutations, the corresponding mutations were generated using the MutateResidue mover in RosettaScripts^75^. Example run files, constraints, options, RosettaScripts XML, and ligand params files are available on GitHub: [https://github.com/hanli-lab/nox-bdh].

## Authors

**Derek Aspacio -** *Department of Chemical and Biomolecular Engineering, University of California, Irvine, Irvine, California 92697, United States*.

**Yulai Zhang -** *Department of Chemical and Biomolecular Engineering, University of California, Irvine, Irvine, California 92697, United States*.

**Youtian Cui -** *Genome Center, University of California, Davis, Davis, California 95616, United States*.

**Edward King -** *Department of Molecular Biology and Biochemistry, University of California, Irvine, Irvine, California 92697, United States*.

**William B. Black -** *Department of Chemical and Biomolecular Engineering, University of California, Irvine, Irvine, California 92697, United States*.

**Sean Perea -** *Department of Chemical and Biomolecular Engineering, University of California, Irvine, Irvine, California 92697, United States*.

**Emma Luu** *-Genome Center, University of California, Davis, Davis, California 95616, United States*.

**Justin B. Siegel** *-Department of Chemistry, University of California, Davis, Davis, California 95616, United States; Genome Center, University of California, Davis, Davis, California 95616, United States; Department of Biochemistry and Molecular Medicine, University of California, Davis, Davis, California 95616, United States*.

## Author contribution

D.A., Y.Z., E.K., and H.L. designed the experiments. Y.C. and E.L performed Rosetta modeling. E.K. and Y.Z. performed rational protein engineering experiments. Y.Z. performed growth-based selection of Nox. D.A. and E.K. bioprospected Bdh homologs. D.A. performed the Michaelis-Menten kinetic experiments for Bdhs. Y.Z. performed the Michaelis-Menten kinetic experiments for Nox. D.A. and S.P. performed cell-free biotransformation experiments. D.A. performed the resting cell biotransformation experiments. W.B. and D.A. reformulated the colorimetric redox ratio assay. D.A. performed the colorimetric redox ratio assays. Y.C., E.L, and J.S. analyzed the modeling results. All authors analyzed the data and wrote the manuscript.

## Acknowledgement

H.L acknowledges support from University of California, Irvine, the national Science Foundation (NSF) (award no. 1847705), and the National Institutes of Health (NIH) (award no. DP2 GM137427), Alfred Sloan research fellowship, and Advanced Research Projects Agency–Energy (ARPA-E) (award no. DE-AR0001508). Y.C., E.L., and J.B.S. acknowledge the funding of the National Institute of Environmental Health Sciences Grant Number: P42ES004699, the National Institutes of Health Grant Number R01 GM 076324-11, and the National Science Foundation Grant Numbers: 1627539, 1805510, 1827246. R.L. acknowledges support from the National Institutes of Health (NIH) (award no. GM130367). D.A acknowledges support from the NSF Graduate Research Fellowship Program (grant no. DGE1839285). S.M acknowledges support from the NSF Graduate Research Fellowship Program (grant no. DGE1839285). The authors acknowledge valuable technical support provided by AAT Bioquest for the redox ratio kits.

## Notes

The authors declare no competing financial interests.

## Supplemental Information

Extended methods; Plasmids and strains used in this study (Table S1); Variants in *Ll* Nox selection (Table S2); Apparent kinetic parameters of engineered enzymes (Table S3); Cofactor preference, databased identifier and source of enzymes used in stereo-upgrading systems (Table S4); Database identifiers and putative enzyme class for other enzymes in bioprospecting campaigns (Table S5); Specific activity of *Ll* Nox I159T and selection variants (Fig. S1); Model of *Ll* Nox WT with NMNH (Fig. S2); *Kp* m-Bdh WT and *Kp m*-Bdh Ortho predicted interactions with NAD^+^ (Fig. S3); Purified protein cycling reactions of NMNH-active variants of Bdh homologs (Fig. S4); *Ser* S-Bdh WT and *Ser* S-Bdh Ortho predicted interactions with NAD^+^ (Fig. S5); Bioprospecting an NADP^+^-active R-Bdh for m-Bdo oxidation (Fig. S6); Specific activity of Nox Ortho for all three cofactors at application condition (Fig. S7).

