## Supporting Information for "Shifting Redox Reaction Equilibria on Demand Using an Orthogonal Redox Cofactor"

*Corresponding author.

**Corresponding Authors**

**Supporting Information**

**A. Extended Methods**

**B. Supplemental Table and Figures**

Table S1 Strains and Plasmids

Table S2. Mutations Observed in *Ll* Nox Variants Obtained From Selection

Table S3. Apparent Kinetic Parameters of Enzymes Engineered for NMN(H)

Table S4. Cofactor Preference of Enzymes Used in Stereo-Upgrading Systems

Table S5. Database Identifiers for Wild Type Sequences of Other Enzymes Used in Bioprospecting Experiments

Figure S1. Specific activity of *Ll* Nox I159T and *Ll* Nox selection variants

Figure S2. Model of *Ll* Nox WT with NMNH

Figure S3. *Kp* m-Bdh WT and *Kp m*-Bdh Ortho predicted interactions with $NAD^+$

Figure S4. Purified protein cycling reactions of NMNH-active variants of Bdh homologs

Figure S5. *Ser* S-Bdh WT and *Ser* S-Bdh Ortho predicted interactions with $NAD^+$

Figure S6. Bioprospecting an $NADP^+$-active R-Bdh for m-Bdo oxidation

Figure S7. Specific activity of Nox Ortho for all three cofactors at application condition

**C. References**

### A. Extended Methods

**Buffers and Selection Media Recipes.** His-Binding Buffer was prepared following the recipes in the His-Spin Protein Miniprep kit (Zymo research) at pH 7.7: 300 mM NaCl, 50 mM sodium phosphate pH 7.7, 10 mM imidazole, and 0.03% Triton X-100. His-Wash Buffer was prepared at pH 7.7: 300 mM NaCl, 50 mM sodium phosphate, 50 mM imidazole, and 0.03% Triton X-100. His-Elution Buffer was prepared at pH 7.7: 300 mM NaCl, 50 mM sodium phosphate, and 250 mM imidazole. 1x phosphate buffered saline (PBS) was prepared as 137 mM NaCl, 2.7 mM KCl, 10 mM sodium phosphate dibasic, 1.8 mM potassium phosphate monobasic, and titrated to pH 7.4. Titration of all buffers was performed with concentrated HCl or 10 M NaOH as appropriate.

M9 Wash Buffer contained 1 mM $MgSO_4$, 0.1 mM $CaCl_2$, BD Difco M9 salts ($Na_2HPO_4$ 6.78 g/L, $KH_2PO_4$ 3g/L, NaCl 0.5 g/L, $NH_4Cl$ 1 g/L), and Sigma Aldrich trace metal mix A5 with Co ($H_3BO_3$ 2860 µg/L, $MnCl_2 \cdot 4H_2O$ 1810 µg/L, $ZnSO_4$ $7H_2O$ 222 µg/L, $Na_2MoO_4$, $2H_2O$ 390 µg/L, $CuSO_4$, $5H_2O$ 79 µg/L, $Co(NO_3)_2 \cdot 6H_2O$ 49 µg/L). M9 Selection Media and Expression Media were prepared as previously described[1]. M9 Selection Media shared the same composition of M9 Wash Buffer with the inclusion of 20 g/L D-glucose, 1 mM nicotinamide, 0.1 mM IPTG, and 0.05% (w/v) L-arabinose, 200 mg/L ampicillin, 50 mg/L kanamycin, and 50 mg/L spectinomycin. For solid media M9 Selection Plates, 15 g/L agar was added in addition to the M9 Selection Media composition. Expression media consisted of 2xYT media supplemented with 20 g/L D-glucose, 1 mM nicotinamide, and antibiotics as above.

**Splicing-by-Overlap Extension PCR.** A standard splicing-by-overlap extension PCR was performed with a KOD Xtreme Hot Start DNA Polymerase PCR reaction mixture (Novagen) at 25 µL total volume and 0.5 µL of each linear DNA fragment with the primers excluded. After a 7-cycle touchdown PCR followed by 5 cycles fixed at the desired annealing temperature, the reactions were removed from the thermocycler. Primers were added to appropriate concentration and a PCR reaction fixed at the annealing temperature was cycled 25-30 times. Once isolated, the vector and insert were circularized by Gibson isothermal assembly[2].

**Protein Expression.** For large scale protein expression cultures, Pyrex #4985 non-baffled shake flasks with phenolic screw-tops were used in place of conical tubes. The flasks were filled to 40 % capacity with 2xYT-200 mg/L ampicillin media and inoculated to 0.07 $OD_{600}$. After induction at 0.5 mM IPTG, the flasks were incubated in a room temperature shaker at ~25 °C at 177 rpm on a 2-inch orbit for ~24 hours. Screw-top flasks were incubated with their caps loose. At harvest, centrifuge bottles were used to pellet the cells. The cell pellets were stored at -80 °C after removal of supernatant.

**Large Scale Lysis by French Pressure Cell Disruption.** The wet cell weight (WCW) of the cell pellet was measured and 1-2 mL of His-binding buffer containing 1 mg/L DNAse and optionally 1 mM AEBSF protease inhibitor was added to the pellets per gram WCW. The cells were fully resuspended on ice. Once fully resuspended the cell suspension was lysed by 3-4 passes through a Thermo Electron French Press with the standard 40K cell at 4 °C. Pressure was maintained at 1000 psig with medium ratio selected and the sample was allowed to exit the outlet at ~15 drops/min. Once cell disruption was complete the lysate was clarified by centrifugation at 4 °C, 21,000 *g* for 15 min at 4 °C. Once pooled, the soluble fraction was ready for purification by immobilized metal affinity chromatography. Generally, a ratio of 100 µL HisPur Ni-NTA resin slurry (Thermo Scientific) equilibrated in His-Binding Buffer was used per mL of clarified lysate.

**Small Scale Lysis by Glass Bead Homogenization.** Each pellet generated from either 20 mL culture grown in the conical tubes or by 40 mL culture distributed from shake flasks was resuspended in His-Binding Buffer containing 1 mM AEBSF protease inhibitor as appropriate. For purification of *Ll* Nox, His-Binding Buffer for resuspension was supplemented with 1 mg/L DNAse and 0.15 mM flavin adenine dinucleotide (FAD). Cell pellets from 20 mL culture were resuspended in 600 μL His-Binding Buffer by vortexing and cell pellets from 40 mL culture were resuspended in 800 μL. Each resuspended cell pellet was transferred to a 2 mL bead beating tube containing 0.5 mL, 0.1 mm diameter glass beads (BioSpec). The bead beating tubes were homogenized in 4 cycles for 35 seconds each at 6.0 m/s in a FastPrep-24 Classic (MP Biomedicals) with 5 min rest on ice between cycles. Once lysed, the samples were clarified of debris and glass beads by centrifugation at 4 °C, 21,000 *g* for 15 min. The soluble fractions of each were transferred to 100 μL of HisPur Ni-NTA resin slurry (Thermo Scientific) equilibrated in His-Binding Buffer.

**Purification of Poly-Histidine Tagged Protein.** All steps performed at 4 °C or on ice. Clarified lysate and Ni-NTA resin were combined. The mixture was allowed to incubate end-over-end mixing (15 rpm) at 4 °C for 15 min to 1 hour. After incubation, the resin and lysate mixture were centrifuged at 4 °C, 700 *g* for 1 min and the supernatant discarded. Six resin bed-volumes (volume of solid resin, BV) of His-Binding Buffer were added to the resin and mixed gently to fully resuspend the resin bed. The entire slurry was transferred to a Thermo Scientific Pierce centrifuge column. The binding buffer was removed by centrifugation at 700 *g* for the Pierce columns. The resin was washed twice with 6 BV His-Wash Buffer. After the resin had been washed twice to remove nonspecific proteins, 1.5-2.5 BVs of His-Elution Buffer was added directly to the resin. After 10 min of gently mixing the final elution was collected in a clean 1.5 mL or 15 mL tubes by centrifugation at 700 *g* for 2 min. The eluant was quantified by Bradford assay relative to a standard curve of bovine serum albumin. Glycerol was added to a final concentration of 20 % and stored in single-use aliquots at -80 °C.

**Salting-Out-Extraction of Butanediol.** An aqueous phase was prepared by mixture of 100 μL of sample with 100 μL 1000 g/L potassium phosphate dibasic solution. 200 μL extraction solution (80%: 20% ethyl acetate: ethanol with 200 mg/L 4-oxoisophorone as an internal standard) was added to the aqueous phase and the extractions were mixed vigorously at maximum speed on an analog vortexer. The phases were separated by centrifugation at 21,000 *g*, 2 min. 100 μL of the organic phase was stored in GC vials with low volume inserts at 4 °C until injection.

**Bioprospecting an R-Bdh with $NADP^+$-Activity.** A cofactor containing crystal structure of *Bs* R-Bdh was approximated by alignment of PDB: 6IE0 (*Bs* R-Bdh) with the cofactor bound crystal structure of greatest pairwise sequence identity in the protein databank (PDB: 1PL6). The atomic coordinates of the $NAD^+$ ligand was transferred to the *Bs* R-Bdh structure, and the 2'-hydroxyl replaced with phosphate in Pymol (Schrodinger). For bioprospecting cofactor specificity reversal homologs, we hypothesized that homologs with mismatched residues at key positions on the loop that would contact the 2'-phosphate of $NADP^+$ would be the best targets. Of the top 2,000 protein-protein BLAST[3] hits of *Bs* R-Bdh against the NCBI nonredundant protein database[4], 891 sequences were found to have at least one mismatch at positions E200, L201, and R205 of *Bs* R-Bdh (Fig. S6A-C). We further filtered the sequences to only include substitution of E200 to glutamine for the potential to create favorable electrostatic interactions with the 2'-phosphate of $NADP^+$. An E200Q hits table of 163 sequences was recorded and sorted by descending pairwise sequence identity. Interestingly, all hits for E200Q also had the substitution L201R. We selected a representative sample of nine sequences throughout the list (Table S1, S4, S5) for codon optimization, DNA synthesis, and plasmid cloning as described in the Main Text.

**Cycling Reaction for R-Bdh Homologs Stereospecificity.** Reactions contained final concentrations of 100 mM potassium phosphate pH 7.5, 1 M NaCl, 200 mM D-glucose, 2mM cofactor, and 5 g/L m-Bdo in a final volume of 450 µL. Each Nox was provided at 11.7 µM and Bdh homologs at 28.1 µM. Protein dilutions were carried out using His-Elution Buffer. Reactions were initiated by addition of 213 µL protein mixture to 237 µL concentrated reaction master mix. Reactions in 15 mL conical tube were incubated, shaking (250 rpm, 45° angle) at 30 °C for 10 hours then sampled as described in the Salting-Out-Extraction of Butanediol section above.

**Reformulation of Commercial Redox Ratio Kits for NMNH/NMN$^+$ quantification.** The Amplite Colorimetric NAD$^+$/NADH Ratio Assay Kit (AAT Bioquest) was used to measure NAD(H) and NAD$^+$, the Amplite Colorimetric NADP$^+$/NADPH Ratio Assay Kit was used to measure NADP(H), and the Amplite Colorimetric NADH Assay Kit with some components from the other kits were reformulated to prepare an NMNH/NMN$^+$ kit. Briefly, a colorimetric method for quantification of NMN$^+$ and total NMN(H) was designed to function in the same way as the commercial kits. In principle, the kits are coupled cycling reactions between an enzyme with strict cofactor specificity and a chromogenic sensor with a maximum absorbance at 460 nm upon reduction by any reduced nicotinamide cofactor. Since only the cofactor recognized by the kit's enzyme is recycled, only one cofactor is turned over repeatedly. Reduced species of the other cofactors, once oxidized by the probe, are not recycled so their contribution to the final absorbance at 460 nm becomes negligible when many turnovers of the target cofactor occur.

To measure only the reduced or oxidized species in a sample, one set of each sample is treated with a vendor supplied (AAT Bioquest) acid extraction solution and heat. In this condition the reduced cofactor is destroyed, the sample is neutralized, and then oxidized cofactor measured. The total pool of a given cofactor is measured by treatment with a vendor supplied control solution (AAT Bioquest) that does not degrade either form. The reduced cofactor is quantified as the difference between the total pool of a given nucleotide and oxidized form of that cofactor. Based on this principle, the Amplite Colorimetric NADH Assay Kit which uses the same probe to directly measure NADH without a recycling enzyme could be reformulated to measure the turnover of NMN(H) if a strictly NMN$^+$-dependent enzyme and substrate were provided. Therefore, we chose to use the previously engineered Gdh Ortho[5] as the recycling enzyme with the NADH probe, assay buffer, and D-glucose. To measure multiple cofactors simultaneously, the samples were treated in parallel following the protocol for each kit. The protocol below is general to all kits as the kits only differ by the assay working solution formulation, appropriate dilution fold to stay within the linear dynamic range, and ratio of sample to master mix volume.

Samples of 75 µL reaction mixture from the SS-Bdo or RR-Bdo stereo-upgrading reactions were aliquoted into microcentrifuge tubes and immediately stored at -80 °C. When all samples were collected, the samples were removed from the freezer to thaw. Immediately once thawed, 150 µL 1x PBS was added to each sample. The samples were mixed by vortexing and centrifuged. To ensure that no enzymes in the reaction mixture can interfere with the assay readout, all protein greater than 10 kDa was removed from each sample by filtration of 200 µL sample through PALL Acroprep Advance 96 well 10K MWCO Omega short tip filtration plate (8034) by centrifugation at 1500 *g* for 10 min, 4 °C. A 96 well plate was placed underneath to collect the protein-free filtrate. Protein-free samples were processed in parallel to the standard curves.

A twelve-point standard curve for the range of 0-2 mM was generated for NAD$^+$, NADP$^+$, and NMN$^+$ in the same buffer composition as the samples. The standard curve was diluted three-fold in 1x PBS. Once it was determined that the NAD$^+$ or NADP$^+$ was entirely oxidized in all NMN(H) containing samples, two separate NMN$^+$ standard curves were prepared containing a constant concentration of either

2 mM $NAD^+$ or 2 mM $NADP^+$. This was to ensure that any change in color at 460 nm attributable to the other cofactors was accounted for because while exceedingly small, some $NAD(P)^+$ activity with Gdh Ortho is possible[5].

Oxidized cofactor sample treatment:

15 µL of protein-free sample was combined with 15 µL of Component D: Extraction Solution using a multichannel pipette and mixed gently by pipetting. Extraction solution will degrade the reduced nucleotides. The samples were then incubated in a thermocycler for 15 min at 37 °C then cooled to 4 °C. Immediately after cooling the samples were removed from the thermocycler and 15 µL of Component E: Neutralization Solution was added and mixed by gentle pipetting with a multichannel. 10 µL aliquots of the processed samples were made for each cofactor determination method ($NAD^+$, $NADP^+$, $NMN^+$). The $NAD^+$ aliquots were diluted 10-fold in 1x PBS, $NADP^+$ 33-fold in 1x PBS, and $NMN^+$ aliquots were diluted 8/3-fold in 1x PBS. 50 µL of diluted $NAD(P)^+$ sample was transferred to fresh PCR tubes that would be used for initiation of the colorimetric cycling reaction. For $NMN^+$, 20 µL of diluted sample was transferred to fresh PCR tubes for initiation of the assay reaction.

Total nucleotide pool (oxidized + reduced single cofactor) and standards sample treatment:
15 µL of protein-free sample was combined with 15 µL of Component F: Control Solution using a multichannel pipette and mixed gently by pipetting. Control solution does not degrade the reduced or oxidized nucleotides. The samples were then incubated in a thermocycler for 15 min at 37 °C then cooled to 4 °C. Immediately after cooling the samples were removed from the thermocycler and 15 µL of Component F: Control Solution was added and mixed by gentle pipetting with a multichannel. 10 µL aliquots of the processed samples were made for each cofactor determination method (NAD(H), NADP(H), NMN(H)). The NAD(H) aliquots were diluted 10-fold in 1x PBS, NADP(H) 33-fold in 1xPBS, and NMN(H) aliquots were diluted 8/3-fold in 1x PBS. 50 µL of diluted NAD(P)(H) sample was transferred to fresh PCR tubes that would be used for initiation of the colorimetric cycling reaction. For NMN(H), 20 µL of diluted sample was transferred to fresh PCR tubes for initiation of the assay reaction.

Assay working solution preparation:

Assay buffers were prepared with the following recipes with NADH or NADPH components from the corresponding commercial kit (AAT Bioquest). The same working solution was used for samples from either sample treatment protocol.

NAD(H) assay buffer was prepared as a bulk working solution and contained 40 µL Component B-II: NADH Probe Buffer per sample, 10 µL Component B-I: NADH Probe per sample, and 1.5 µL Component A: NAD/NADH Recycling Enzyme Mix per sample.

NADP(H) assay buffer was prepared as a bulk working solution and contained 40 µL Component B-II: NADPH Probe Buffer per sample, 10 µL Component B-I: NADPH Probe per sample, and 1.5 µL Component A: NADP/NADPH Recycling Enzyme Mix per sample.

NMN(H) assay buffer was prepared as a bulk working solution using components from the Amplite Colorimetric NADH Assay Kit and contained the following: 40 µL Component B: Assay Buffer per sample, 10 µL Component A: NADH Probe per sample, 10 µL of 200 g/L D-glucose per sample, 5 µL 1x PBS per sample, and 15 µL 67 µM Gdh Ortho per sample. Gdh Ortho was stored at -80 °C in high concentration as single use aliquots prior to addition to the working solution, dilution to 67 µM from stock was done with his-elution buffer as the diluent.

Reaction initiation:

For NAD(P)(H) determination, 50 µL of NAD(P)(H) assay working solution was distributed to clean PCR tubes. For NMN(H) assay working solution, 80 µL was distributed to clean PCR tubes. To initiate the colorimetric recycling reaction assay, working solution was transferred to 20 µL treated samples by multichannel pipette, mixed gently by pipetting, and transferred to a 96 well visible plate at room temperature. The plate was incubated at room temperature without agitation or exposure to light for 1-3 hours then the endpoint absorbance at 460 nm was recorded.

Data Analysis:

The 0 mM nucleotide standard for each nucleotide was used as the blank for all samples of that assay type. The equation of the best fit line to the corresponding standard curve was used to quantify the concentration of the target nucleotide in the samples. Reduced cofactor was calculated as [total pool of a single nucleotide] - [Oxidized]. Three replicate samples were processed per condition, per cofactor, and per treatment type (oxidized or total pool). Error of the total pool and oxidized species concentrations is calculated as the standard deviation of three independent replicates. Error in the concentration of reduced species and redox ratio was calculated by propagation of error. Reduced cofactor concentrations below a defined limit of quantification were set at the limit of quantification based on confidence interval as described in the Limit of Reduced Cofactor Quantification section of the Main Text.

### B. Supplemental Table and Figures

**Table S1. Strains and Plasmids**

| **Strains** | **Description** | **Reference** |
|---|---|---|
| XL-1 blue | *E. coli* Cloning strain | Stratagene |
| BL21 (DE3) | *E. coli* Protein expression strain | Invitrogen |
| BW25113 | *E. coli* $F^-$, *lacI*$^q$ *rrnB*$_{T14}$ Δ*lacZ*$_{WJ1}$ *hsdR514* Δ*araBAD*$_{AH33}$ Δ*rhaBAD*$_{LD78}$ | Datsenko *et al.* 2000[6] |
| MX102 | BW25113 *ΔpncC Δpgi Δzwf ΔgntK::Kan*$^R$ | Black *et al*. 2020[5] |
| MX102 $R^0$ | BW25113 *ΔpncC Δpgi Δzwf ΔgntK* | This study |
| MX502 | BW25113 *ΔpncC Δpgi Δzwf ΔnadR Δgnd* + pLM106 + pLS502 | King *et al*. 2022[1] |

| **Plasmids** | **Description** | **Reference** |
|---|---|---|
| pCP20 | Temperature-inducible yeast Flp recombinase gene controlled by λ*c*I*ts857* in a temperature-sensitive replicon | Datsenko *et al.* 2000[6] |
| pDA063 | $P_{LlacO1}$:: *Bs bdhA* (*Bs* R-Bdh), ColE1 *ori,* Amp$^R$, N-terminal 6x His tag | This study |
| pDA091 | $P_{LlacO1}$:: *Ecl dar* Y34Q-A87K-M189T, ColE1 *ori,* Amp$^R$, N-terminal 6x His tag | This study |
| pDA092 | $P_{LlacO1}$:: *Kp dar* (*Kp* m-Bdh WT), ColE1 *ori,* Amp$^R$, N-terminal 6x His tag | This study |
| pDA105 | $P_{LlacO1}$:: *As bdh,* ColE1 *ori,* Amp$^R$, N-terminal 6x His tag | This study |
| pDA106 | $P_{LlacO1}$:: *Cr bdh,* ColE1 *ori,* Amp$^R$, N-terminal 6x His tag | This study |
| pDA107 | $P_{LlacO1}$:: *Ca bdh,* ColE1 *ori,* Amp$^R$, N-terminal 6x His tag | This study |
| pDA108 | $P_{LlacO1}$:: *Pb bdh,* ColE1 *ori,* Amp$^R$, N-terminal 6x His tag | This study |
| pDA109 | $P_{LlacO1}$:: *Cbo bdh,* ColE1 *ori,* Amp$^R$, N-terminal 6x His tag | This study |
| pDA110 | $P_{LlacO1}$:: *Km bdh,* ColE1 *ori,* Amp$^R$, N-terminal 6x His tag | This study |
| pDA111 | $P_{LlacO1}$:: *Cs bdh* (*Cs* R-Bdh), ColE1 *ori,* Amp$^R$, N-terminal 6x His tag | This study |
| pDA112 | $P_{LlacO1}$:: *Cbu bdh,* ColE1 *ori,* Amp$^R$, N-terminal 6x His tag | This study |
| pDA113 | $P_{LlacO1}$:: *Zp bdh,* ColE1 *ori,* Amp$^R$, N-terminal 6x His tag | This study |
| pDA115 | $P_{LlacO1}$:: *Ser bdh*2 (S-Bdh WT), ColE1 *ori,* Amp$^R$, N-terminal 6x His tag | This study |

| | | |
|---|---|---|
| pDA129 | $P_{LlacO1}$:: *Ser bdh*2 L39Q-A92K-M194T – *Bs gdh* S17E-Y39Q-A93K-I195R, ColE1 *ori,* Amp$^R$, N-terminal 6x His tag | This study |
| pDA130 | $P_{LlacO1}$:: *Ser bdh*2 – *Bs gdh,* ColE1 *ori,* Amp$^R$, N-terminal 6x His tag | This study |
| pDA131 | $P_{BAD}$:: *Lb nox* – *Bs bdhA,* RSF1030 *ori,* Spec$^R$ | This study |
| pDA162 | $P_{BAD}$:: Tp *nox* – *Cs bdh,* RSF1030 *ori,* Spec$^R$ | This study |
| pEK101 | $P_{LlacO1}$:: *Bs gdh,* ColE1 *ori,* Amp$^R$, N-terminal 6x His tag | Black et al. 2020[5] |
| pEK370 | $P_{LlacO1}$:: *Kp budC* Y34Q-A87K-M189T, ColE1 *ori,* Amp$^R$*,* N-terminal 6x His tag | This study |
| pEK463 | $P_{LlacO1}$:: *Kp dar* Y34Q-A87K-M189T (m-Bdh Ortho), ColE1 *ori,* Amp$^R$*,* N-terminal 6x His tag | This study |
| pEK484 | $P_{LlacO1}$:: *Ka budC* Y34Q-A87K-M189T, ColE1 *ori,* Amp$^R$*,* N-terminal 6x His tag | This study |
| pEK485 | $P_{LlacO1}$:: *Ka dar* Y34Q-A87K-M189T, ColE1 *ori,* Amp$^R$*,* N-terminal 6x His tag | This study |
| pEK486 | $P_{LlacO1}$:: *Ko budC* Y34Q-A87K-M189T, ColE1 *ori,* Amp$^R$*,* N-terminal 6x His tag | This study |
| pEK488 | $P_{LlacO1}$:: *Ser bdh2* L39Q-A92K-M194T (*Ser* S-Bdh Ortho), ColE1 *ori,* Amp$^R$, N-terminal 6x His tag | This study |
| pLZ216 | $P_{LlacO1}$:: *Bs gdh* S17E-Y39Q-A93K-I195R (Gdh Ortho)*,* ColE1 *ori,* Amp$^R$, N-terminal 6x His tag | Black et al. 2020[5] |
| pLS401 | $P_{LlacO1}$:: *Lb nox,* ColE1 *ori,* Amp$^R$, N-terminal 6x His tag | Maxel et al. 2021[7] |
| pLS402 | $P_{LlacO1}$:: Tp *nox* (*Lb nox* G159A-D177A-A178R-M179S-P184R)*,* ColE1 *ori,* Amp$^R$, N-terminal 6x His tag | Maxel et al. 2021[7] |
| pYZ10 | $P_{LlacO1}$:: *Ll nox,* ColE1 *ori,* Amp$^R$, N-terminal 6x His tag | This study |
| pYZ11 | $P_{LlacO1}$:: *Ll nox* I159T*,* ColE1 *ori,* Amp$^R$, N-terminal 6x His tag | This study |
| pYZ100 | $P_{LlacO1}$:: *Ll nox* I159T-D178X-A179X-I243X*,* ColE1 *ori,* Amp$^R$, N-terminal 6x His tag | This study |
| pYZ35 | $P_{LlacO1}$:: *Ll nox* I159T-D178N-A179F-I243E (Nox Ortho)*,* ColE1 *ori,* Amp$^R$, N-terminal 6x His tag | This study |
| pSM10 | $P_{LlacO1}$:: *Zm glf,* P15A *ori,* Kan$^R$ | This study |
| pLS501 | $P_{LlacO1}$:: *Pp xenA* D116E*,* ColE1 *ori,* Amp$^R$, N-terminal 6x His tag | King *et al*. 2022[1] |
| pLS502 | $P_{LlacO1}$:: *Ft nadE* – *Ft nadV* – *Zm glf,* P15A *ori*, Kan$^R$ | King *et al*. 2022[1] |
| pLM106 | $P_{BAD}$:: *Bs gdh* S17E-Y39Q-A93K-I195R *,* RSF1030 *ori,* Spec$^R$ | King *et al*. 2022[1] |

Abbreviations indicate source organism of gene: *Bs*, *Bacillus subtilis; Ecl, Enterobacter cloacae* ssp. *dissolvens* SDM*; Kp, Klebsiella pneumoniae; As*, *Atopobium* sp.; *Cr*, *Clostridium ragsdalei*; *Ca*, Clostridium amylolyticum; *Pb*, *Paraclostridium bifermentans*; *Cbo*, *Clostridium botulinum*; *Km*, *Khelaifiella massiliensis*; *Cs*, *Clostridium saccharoperbutylacetonicum*; *Cbu*, *Clostridium butyricum*; *Zp*, *Zymobacter palmae*; Lb, *Lactobacillus brevis* strain ATCC 367; Tp, triphosphopyridine (engineered from *Lb*); *Zm, Zymomonas mobilis*; *Ll, Lactobacillus lactis*; *Ft, Francisella tularensis*; *Ser, Serratia* sp. AS13; *Ka, Klebsiella aerogenes* KCTC 2190; *Ko, Klebsiella oxytoca* KCTC 1686.

Accession numbers of primary genes in these systems included in Table S4.

Accession numbers of bioprospected genes included in Table S5.

**Table S2. Mutations Observed in *Ll* Nox Variants Obtained from Library Selection.**

| Variant | Asp 178 | | Ala 179 | | Ile 243 | |
|---|---|---|---|---|---|---|
| | Codon | Amino Acid | Codon | Amino Acid | Codon | Amino Acid |
| WT | GAT | D | GCG | A | ATT | I |
| LL-1 | TTG | L | GCT | A | AAG | K |
| LL-2 | TAG | Stop | AAG | K | CTG | L |
| LL-3 | CCT | P | AAG | K | AGG | R |
| LL-4 | CCG | P | GAG | E | AAT | N |
| LL-5 | TTG | L | GAT | D | GGT | G |
| LL-6 | GTG | V | GCG | A | CCG | P |
| LL-7 | TGG | W | CAT | H | GCG | A |
| LL-8 | CTT | L | GCG | A | CGG | R |
| Nox Ortho | AAT | N | TTT | F | GAG | E |
| LL-10 | CCT | P | TGT | C | TTG | L |
| LL-11 | CCG | P | CTG | L | CAT | H |
| LL-12 | GGG | G | GTG | V | GAT | D |
| LL-13 | AAG | K | CAT | H | TTT | F |
| LL-14 | GGT | G | TAT | Y | CGG | R |
| LL-15 | ATG | M | GTG | V | AGG | R |
| LL-16 | GTT | V | ACG | T | CGT | R |
| LL-17 | TTG | L | GTC | V | TCG | S |
| LL-18 | TTG | L | CCT | P | AGG | R |

**Table S3. Apparent Kinetic Parameters of Enzymes Engineered for NMN(H).**

| Enzyme | Variant | Cofactor | $K_m$ (mM) | $k_{cat}$ ($s^{-1}$) | $k_{cat}/K_m$ ($mM^{-1}s^{-1}$) |
|---|---|---|---|---|---|
| *Kp* m-Bdh | WT | $NAD^+$ | 0.20 ± 0.04 | 69 ± 1 | 355 ± 70 |
| | | $NADP^+$ | 2.6 ± 0.1 | 0.43 ± 0.03 | 0.16 ± 0.01 |
| | | $NMN^+$ | 7.2 ± 0.4 | 0.12 ± 0.01 | 0.016 ± 0.001 |
| | Ortho Y34Q-A87K-M189T | $NAD^+$ | 6.6 ± 0.5 | 0.015 ± 0.001 | 0.0022 ± 0.0001 |
| | | $NADP^+$ | 2.9 ± 0.2 | 0.0067 ± 0.0004 | 0.0023 ± 0.0001 |
| | | $NMN^+$ | 6.7 ± 0.3 | 2.0 ± 0.3 | 0.30 ± 0.03 |
| *Ser* S-Bdh | WT | $NAD^+$ | 0.22 ± 0.03 | 20 ± 1 | 96 ± 20 |
| | | $NADP^+$ | 1.5 ± 0.2 | 0.20 ± 0.01 | 0.13 ± 0.01 |
| | | $NMN^+$ | 5.6 ± 0.3 | 0.043 ± 0.004 | 0.0079 ± 0.001 |
| | Ortho L39Q-A92K- M194T | $NAD^+$ | 1.6 ± 0.1 | 0.019 ± 0.001 | 0.012 ± 0.001 |
| | | $NADP^+$ | 1.4 ± 0.1 | 0.0020 ± 0.0001 | 0.0014 ± 0.0001 |
| | | $NMN^+$ | 4.3 ± 0.4 | 0.38 ± 0.04 | 0.086 ± 0.002 |
| *Ll* Nox | WT | NADH | 0.045 ± 0.002 | 92 ± 2 | 2000 ± 90 |
| | | NADPH | – | – | 17 ± 1 |
| | | NMNH | – | – | 0.22 ± 0.001 |
| | Ortho I159T-D178N-A179F-I243E | NADH | – | – | 20 ± 0.2 |
| | | NADPH | – | – | 20 ± 1 |
| | | NMNH | – | – | 55 ± 4 |

"–" denotes no data if the apparent $K_m$ and $k_{cat}$ could not be determined precisely. Apparent catalytic efficiency in these cases were instead calculated based on a linear fit to a modified Michaelis-Menten equation under the assumption $K_m \gg$ cofactor concentration. WT, wild type. See Main Text for detailed derivation of equations used.

**Table S4. Cofactor Preference and Database Identifiers for Primary Enzymes in Stereo-Upgrading Systems.**

| Enzyme | UniProt Accession | Cofactor Preference | Source |
|---|---|---|---|
| *Ser* Bdh2 | A0A7U3Z3T2 | NAD(H) | Zhang et al. 2016[8] |
| *Kp* Dar | D7RP28 | NAD(H) | Li et al. 2018[9] |
| *Bs* BdhA | O34788 | NAD(H) | Yan et al. 2009[10] |
| *Cs* Bdh | M1MWX5 | NAD(H) and NADP(H) | This study |
| *Bs* Gdh | P12310 | $NAD^+$ and $NADP^+$ | Black et al. 2020[5] |
| *Ll* Nox | A2RIB7 | NADH | Lopez et al. 2001[11] |
| *Lb* Nox | Q03Q85 | NADH | Titov et al. 2016[12] |
| Tp Nox | Mutated variant of *Lb* Nox[a] | NADPH | Cracan et al. 2017[13] |

[a] Tp Nox (*Lb* Nox G159A-D177A-A178R-M179S-P184R)

**Table S5. Database Identifiers for Wild Type Sequences of Other Enzymes Used in Bioprospecting Experiments.**

| Enzyme | Accession | Enzyme Class |
|---|---|---|
| *Ecl* Dar | GenBank\| JN035909 | m-Bdh |
| *Kp* BudC | PDB\| 1GEG | m-Bdh |
| *Ka* BudC | GenBank\| AEG98409 | m-Bdh |
| *Ka* Dar | NCBI Ref\| WP_015366942 | m-Bdh |
| *Ko* BudC | GenBank\| AEX06195 | m-Bdh |
| *As* Bdh | GenBank\| NLH91458 | R-Bdh |
| *Cr* Bdh | GenBank\| AEI90716 | R-Bdh |
| *Ca* Bdh | NCBI Ref\| WP_073006451 | R-Bdh |
| *Pb* Bdh | NCBI Ref\| WP_148550966 | R-Bdh |
| *Cbo* Bdh | NCBI Ref\| WP_075141790 | R-Bdh |
| *Km* Bdh | NCBI Ref\| WP_102401827 | R-Bdh |
| *Cbu* Bdh | NCBI Ref\| WP_104675707 | R-Bdh |
| *Zp* Bdh | NCBI Ref\| WP_027704711 | R-Bdh |

PDB, Protein Data Bank; NCBI Ref, NCBI Reference Sequence Database; m-Bdh, Enzyme Class 1.1.1.B20 (meso-2,3-butanediol dehydrogenase); S-Bdh, Enzyme Class 1.1.1.76 ((S,S)-butanediol dehydrogenase); R-Bdh, Enzyme Class 1.1.1.4 ((R,R)-butanediol dehydrogenase). See Table S1 for genus abbreviations.

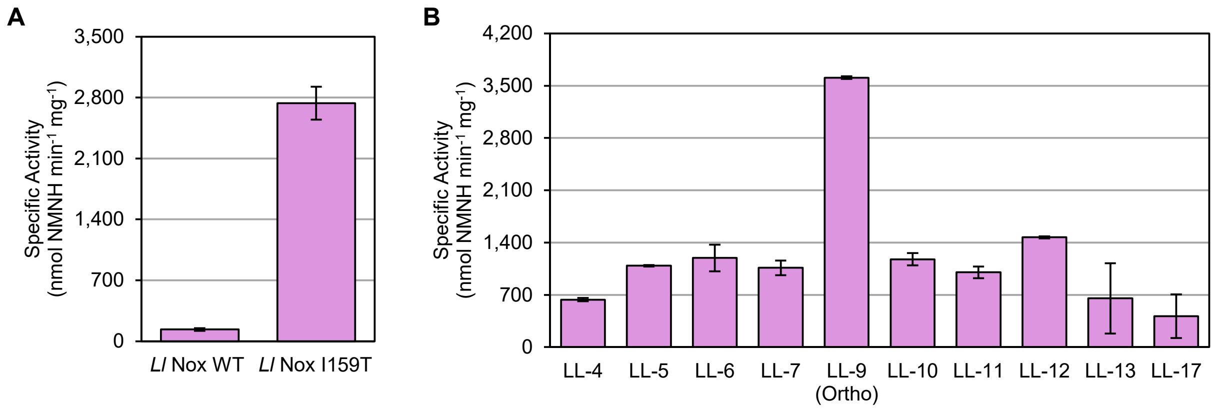


**Figure S1. Specific activity of *Ll* Nox I159T and *Ll* Nox selection variants. (A)** Comparison of *Ll* Nox WT and *Ll* Nox I159T. **(B)** Specific activity toward NMNH for candidate *Ll* Nox variants from selection. LL-9 (Ortho), renamed Nox Ortho, showed ~30-fold increase in NMNH activity compared to WT. Data are presented as mean of two replicates (n=2) ± one standard deviation.

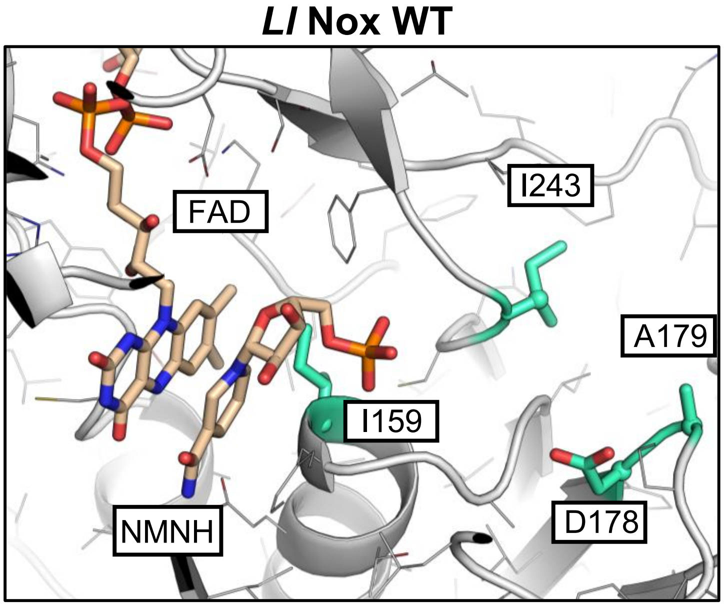


**Figure S2. Model of *Ll* Nox WT with NMNH.** NMNH atomic coordinates are taken as the corresponding position of the NMN-moiety of an NADH bound in homologous structures.

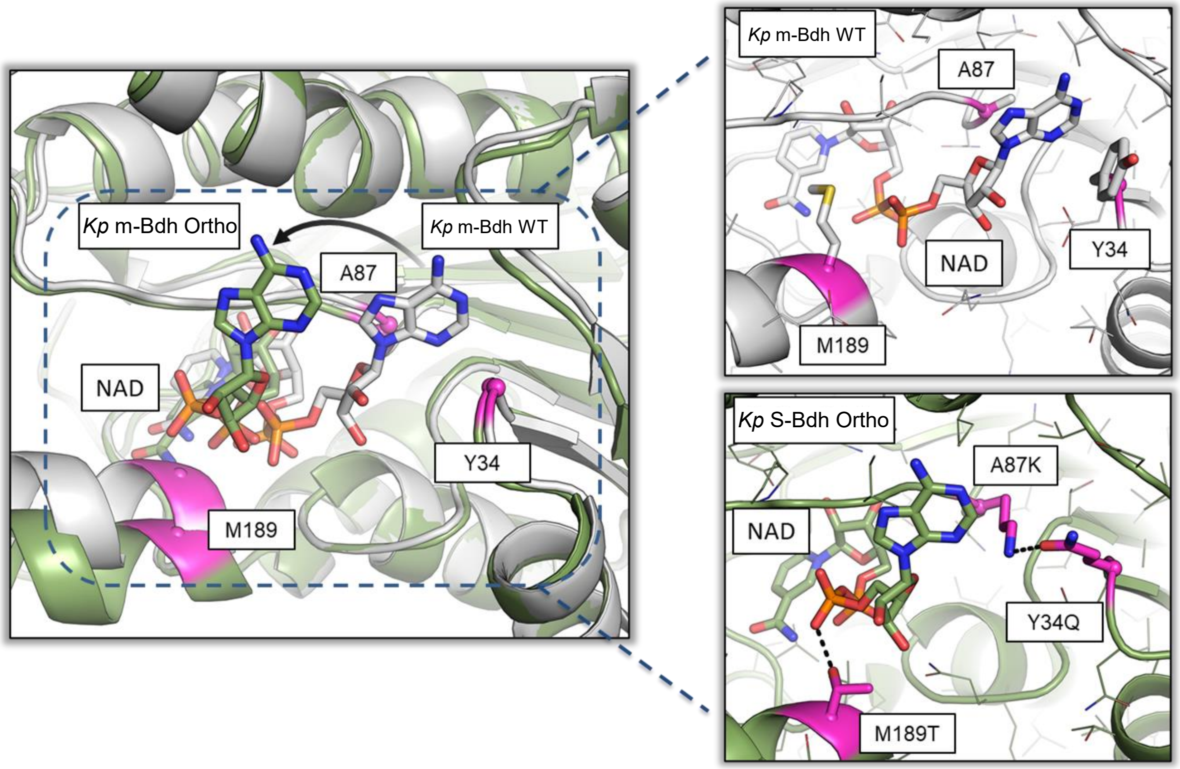


**Figure S3. *Kp* m-Bdh WT and *Kp* m-Bdh Ortho predicted interactions with NAD$^+$.** Compared to *Kp* m-Bdh WT, *Kp* m-Bdh Ortho adopts a nonnatural, nonproductive NAD$^+$ conformer. Despite the polar interaction introduced by M189T, the mutations of A87K and Y34Q resulted in malicious effect of NAD$^+$ binding by forming a novel hydrogen bond interaction that exclude the adenine group of NAD$^+$ while removing the favorable hydrophobic interaction between the cofactor and side chains of A87 and Y34

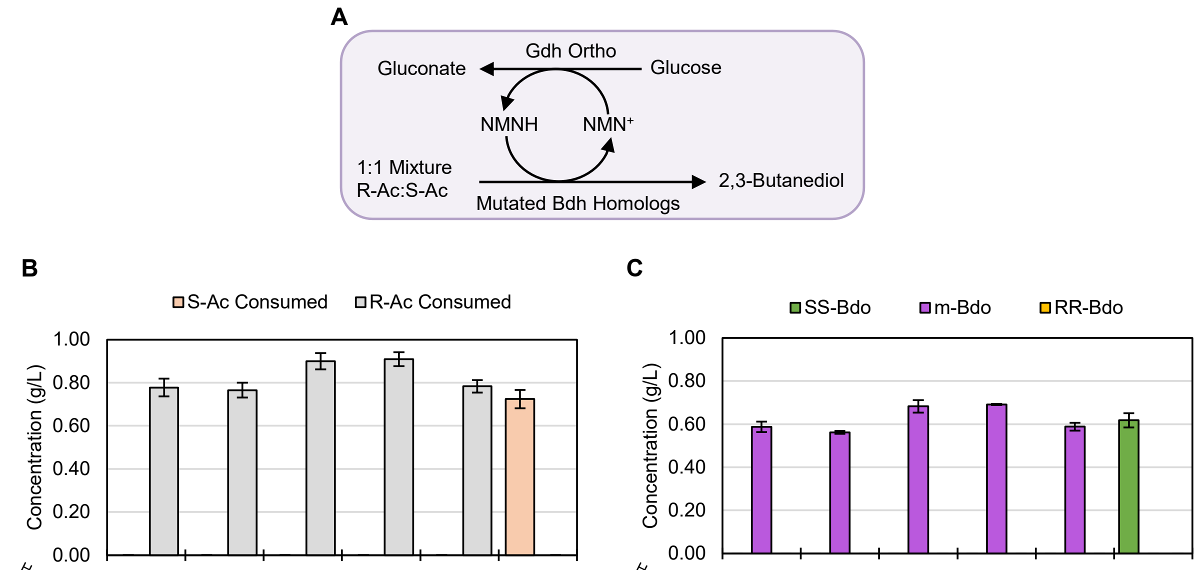


| Homolog / *Kp* m-Bdh sites | *Kp* m-Bdh Ortho | *Ka* BudC Ortho | *Ka* Dar Ortho | *Kp* BudC Ortho | *Ec* Dar Ortho | *Ser* Bdh Ortho |
|---|---|---|---|---|---|---|
| **Y34** | Y34Q | Y34Q | Y34Q | Y34Q | Y34Q | Y39Q |
| **A87** | A87K | A87K | A87K | A87K | A87K | A92K |
| **M189** | M189T | M189T | M189T | M189T | M189T | M194T |
| **Substrate Consumed** | **R-Ac** | **R-Ac** | **R-Ac** | **R-Ac** | **R-Ac** | **S-Ac** |

| Homolog / *Kp* m-Bdh sites | *Kp* m-Bdh Ortho | *Ka* BudC Ortho | *Ka* Dar Ortho | *Kp* BudC Ortho | *Ec* Dar Ortho | *Ser* Bdh Ortho |
|---|---|---|---|---|---|---|
| **Y34** | Y34Q | Y34Q | Y34Q | Y34Q | Y34Q | Y39Q |
| **A87** | A87K | A87K | A87K | A87K | A87K | A92K |
| **M189** | M189T | M189T | M189T | M189T | M189T | M194T |
| **Product Formed** | **m-Bdo** | **m-Bdo** | **m-Bdo** | **m-Bdo** | **m-Bdo** | **SS-Bdo** |

**Figure S4. Purified protein cycling reactions of NMNH-active S-Bdh homologs**. **(A)** Design of the cycling reactions. Racemic acetoin (1:1 mixture of enantiomers) is converted into 2,3-butanediol stereoisomers to test the stereospecificity of the engineered Bdh homologs. Reactions contain 1 M NaCl, 100 mM potassium phosphate pH 7.5, 200 mM D-glucose, 2 mM $NMN^+$, 5 g/L racemic acetoin, Gdh Ortho, and one of the Bdh homolog variants with the corresponding mutations. **(B)** Concentration of each acetoin stereoisomer consumed after 4 hours of cycling reaction. Since only one acetoin isomer decreased in each reaction, consumption is calculated as difference in each acetoin isomer's concentration where error bars represent standard deviation of the difference calculated by propagation of error. Based on observed acetoin isomer consumption, the substrate specificity was summarized for each mutant in the table. **(C)** Concentration of each butanediol stereoisomer formed in the same reaction. Only the mutant of *Ser* Bdh forms detectable SS-Bdo. Both *Kp* m-Bdh Ortho and *Ser* S-Bdh Ortho are stereospecific for (S)-installation because the stereoisomer installed for all enzymes tested is an S-chiral center. Values are the average of three replicates (n=3) with error bars representing one standard deviation.

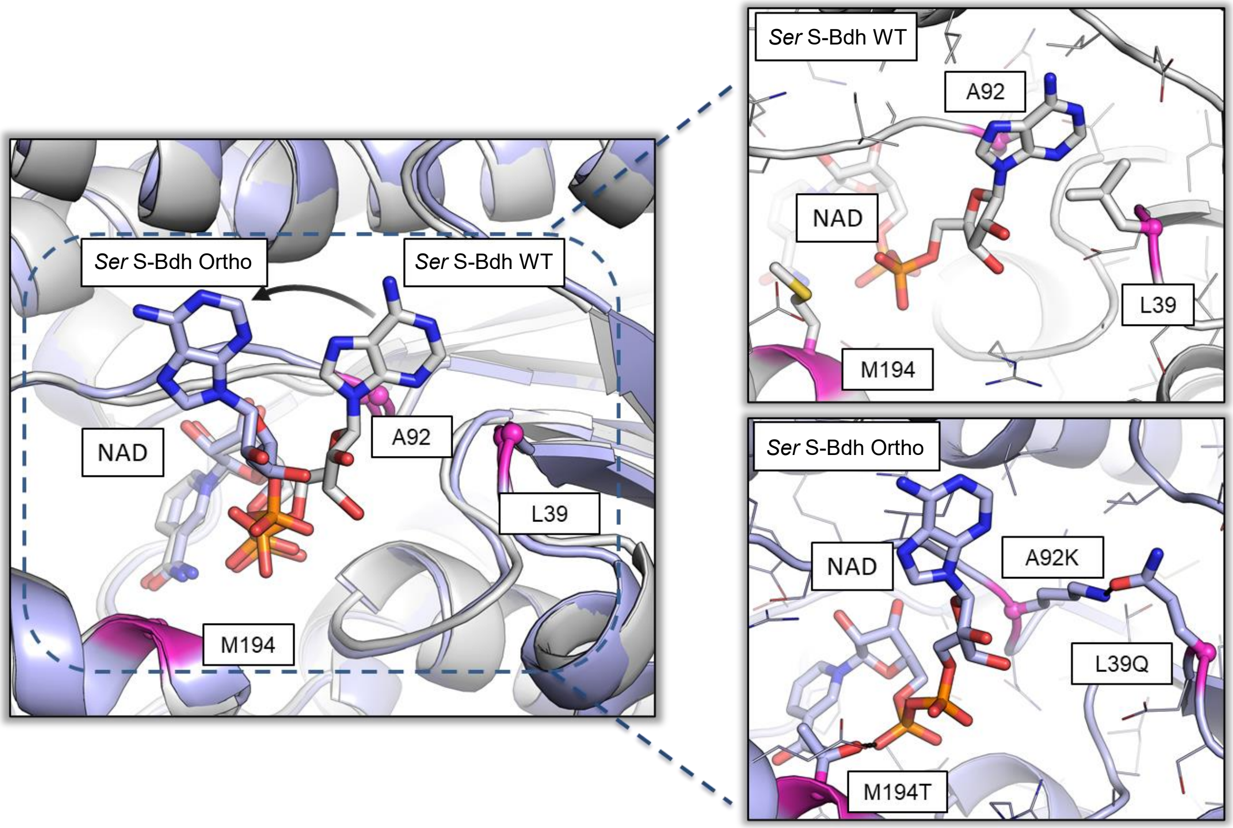


**Figure S5. *Ser* S-Bdh WT and *Ser* S-Bdh Ortho predicted interactions with $NAD^+$.** Similar to the models of *Kp* m-Bdh WT and *Kp* m-Bdh Ortho docked with $NAD^+$ in Fig. S4, *Ser* S-Bdh Ortho is predicted to adopt an unnatural conformer that is not favored by *Ser* S-Bdh WT, leading to the significantly lower catalytic efficiency with $NAD^+$.

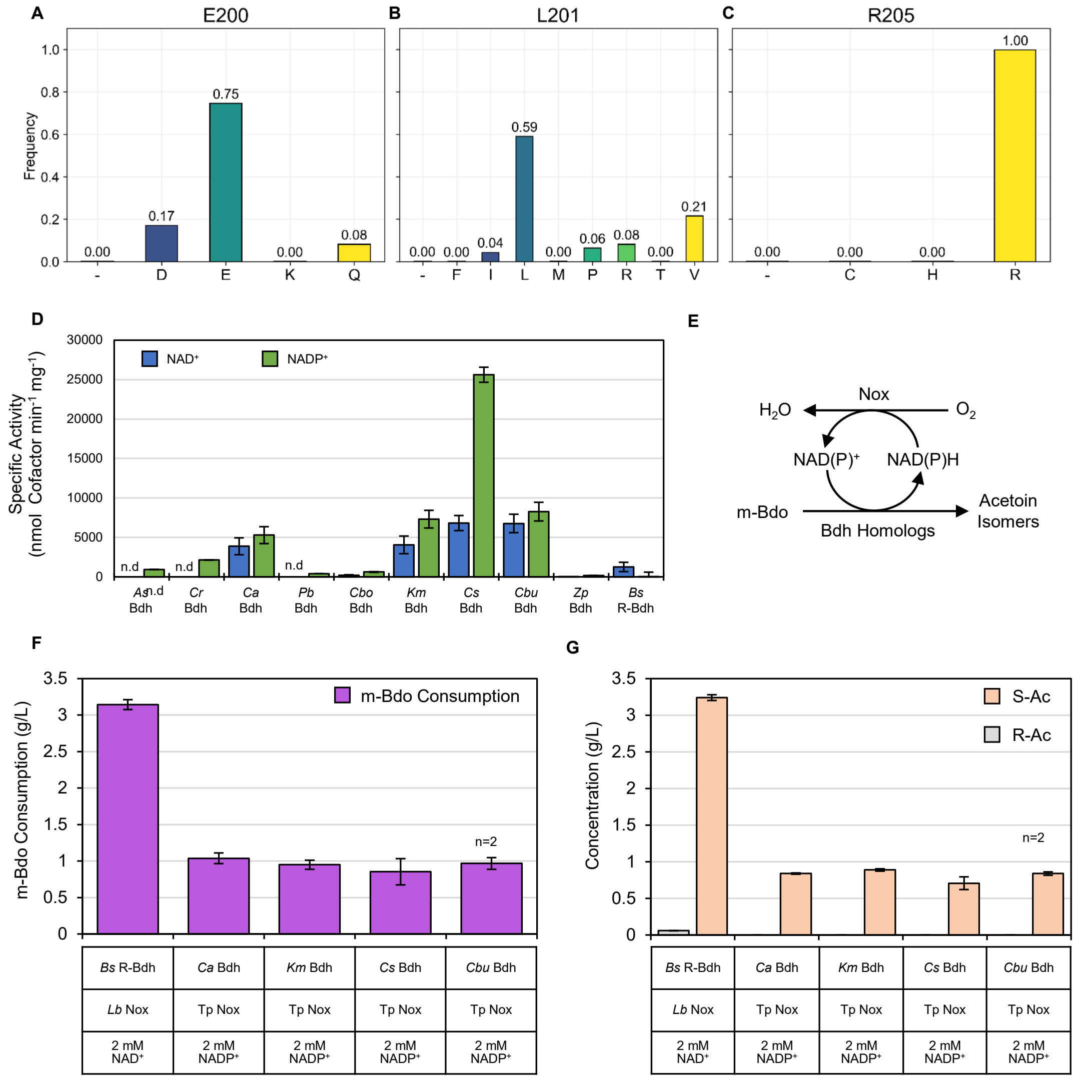


**Figure S6. Bioprospecting an NADP$^+$-active R-Bdh for m-Bdo oxidation. (A-C)** BLAST hits of *Bs* R-Bdh homologs in the NCBI nonredundant protein database[3,4]. Amino acid frequency distributions at **(A)** E200, **(B)** L201, or **(C)** R205 (*Bs* R-Bdh numbering) are shown. Frequency is normalized to 2000 sequences. **(D)** Specific activity of *Bs* R-Bdh homologs with NAD$^+$ and NADP$^+$. Values are the average of two replicates for NAD$^+$ and three replicates for NADP$^+$ with error bars representing one standard deviation. "n.d." denotes that no data was collected for that protein with NAD$^+$. **(E)** Setup of the oxidation cycling assay to determine R-Bdh homolog activity and stereospecificity. R-Bdh activity generates S-Ac, S-Bdh activity generates R-Ac based on the chiral center destroyed. Four *Bs* Bdh homologs were tested for their stereospecificity in the removal of (R)-chiral centers against *Bs* R-Bdh control reaction. **(F)** m-Bdo consumption relative to a no cofactor, no enzyme control reaction. **(G)** Acetoin isomer concentrations. R-Ac is detected in the control reaction for *Bs* R-Bdh which suggests its stereospecificity is not complete. All four bioprospected R-Bdhs tested are stereospecific for the R-chiral center. **(F, G)** Values are the average of three replicates except with *Cbu* Bdh which was performed in duplicate (n=2). Error bars represent one standard deviation.

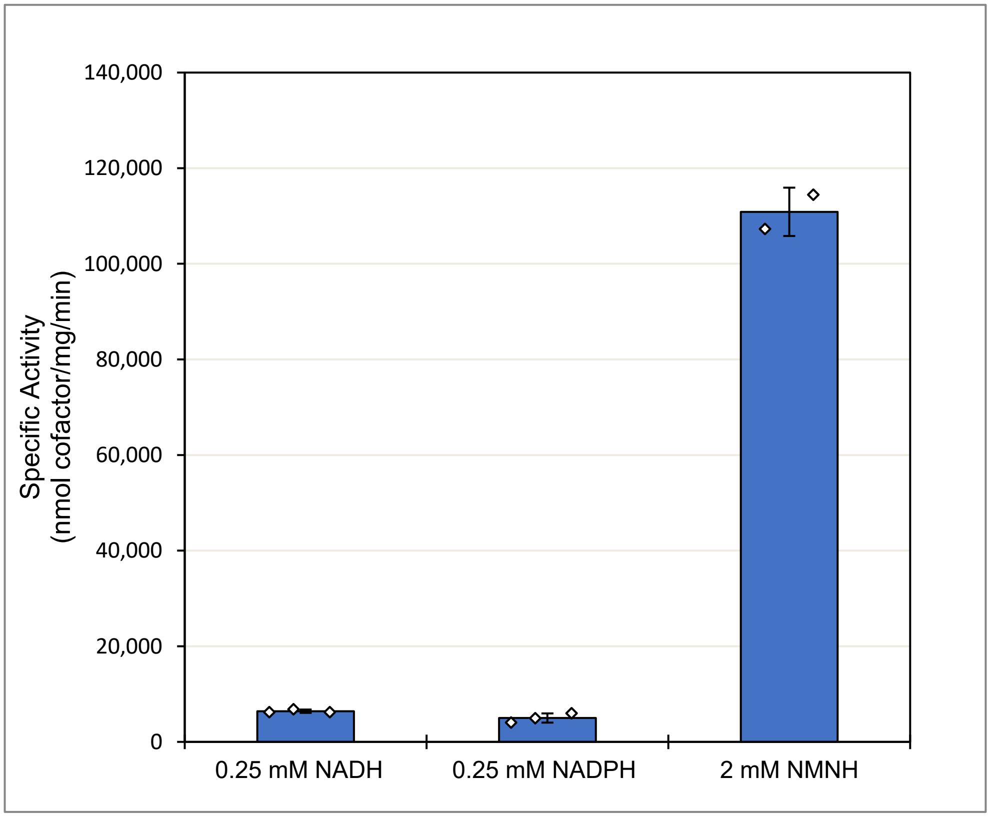


**Figure S7. Specific activity of Nox Ortho for all three cofactors at application condition.** Nox Ortho is significantly more active for NMNH than NAD(P)H in the condition used for the in vitro RR-Bdo production cascade. Assay at 37 °C and 50 mM Tris-Cl at pH 7.0 with listed concentration of reduced cofactor provided.

11. Lopez de Felipe, F.; Hugenholtz, J. Purification and Characterisation of the Water Forming

NADH-Oxidase from Lactococcus Lactis. *Int. Dairy J.* **2001**, *11* (1–2), 37–44. https://doi.org/10.1016/S0958-6946(01)00031-0.

12. Titov, D. V.; Cracan, V.; Goodman, R. P.; Peng, J.; Grabarek, Z.; Mootha, V. K. Complementation of Mitochondrial Electron Transport Chain by Manipulation of the NAD+/NADH Ratio. *Science* **2016**, *352* (6282), 231–235. https://doi.org/10.1126/science.aad4017.

13. Cracan, V.; Titov, D. V.; Shen, H.; Grabarek, Z.; Mootha, V. K. A Genetically Encoded Tool for Manipulation of NADP+/NADPH in Living Cells. *Nat. Chem. Biol.* **2017**, *13* (10), 1088–1095. https://doi.org/10.1038/nchembio.2454.
